# Plasmid copy number variation impacts pathogenicity and quantification of *Curtobacterium flaccumfaciens* pv. *flaccumfaciens* infecting mung bean

**DOI:** 10.1101/2025.01.22.634406

**Authors:** Ahmed Saad, Niloofar Vaghefi, Barsha Poudel, Anthony J. Young, Lisa A. Kelly, Noel L. Knight

## Abstract

In bacteria, plasmids can confer the ability to cause disease. Although they can potentially vary in copy number, little has been reported on the dynamics of plasmids in plant pathogenic bacteria. Pathogenicity of the bacterium *Curtobacterium flaccumfaciens* pv. *flaccumfaciens* (*Cff*), which causes foliar disease on leguminous crops, including mung bean (*Vigna radiata*), has previously been linked to a plasmid. This study explored the variation in plasmid copy number among a genetically diverse collection of 25 *Cff* isolates using purposely designed quantitative PCR assays for chromosomal and plasmid DNA targets. Pathogenicity and virulence of six *Cff* isolates, including one plasmid-free isolate, were assessed on the susceptible mung bean cultivar Opal-AU using visual symptoms, trifoliate dry weights and *Cff* DNA quantities following stem inoculation. Plasmid copy numbers varied significantly (*p* <0.001) among plasmid-carrying isolates, ranging from 1.37 to 2.74. For subsequent quantification *in planta*, the single-copy chromosomal PCR target gene, gyrase b, was selected. Mung beans inoculated with plasmid-carrying isolates exhibited significant variation in visual symptoms (*p* <0.001), while trifoliate dry weights and *Cff* DNA quantities did not significantly differ. The treatment using a plasmid-free isolate was not significantly different from the negative control for each of the disease traits. Knowledge of plasmid dynamics in *Cff* populations lays a foundation for improved understanding of the inheritance and impact of plasmid-related traits. The quantification assays will be useful for monitoring *Cff* populations and the demonstrated variation in pathogenicity and virulence can assist efforts to breed host plant germplasm with reduced susceptibility to *Cff*.

## 1 INTRODUCTION

Bacterial diseases of plants are a persistent and evolving threat to crop production (Sharma et al. 2022). Disease management strategies rely on tactics to reduce pathogen populations in the environment and genetic improvement programs to breed less susceptible crop varieties. The effectiveness of these strategies requires comprehensive knowledge of the factors affecting bacterial pathogenicity and their interactions with the host and environment. The genetic features conferring pathogenicity may be located outside of the bacterial chromosome, on self-replicating, extra-chromosomal DNA molecules known as plasmids (Clark et al. 2019). The dynamics of plasmids among isolates of the same species has had limited investigation, with examples of the presence or absence of a plasmid determining pathogenic or beneficial impacts (Bultreys & Gheysen 2023; Melnyk & Haney 2017), the proficiency of the plasmid host species affecting the maintenance of the plasmid in a population (Kottara et al. 2021), and the copy number of plasmids playing a role in the transmission to new bacterial generations (Hernández-Beltrán et al. 2021; Ilhan et al. 2018).

*Curtobacterium flaccumfaciens* pv. *flaccumfaciens* (*Cff*), a gram-positive, vascular pathogen that systemically colonises leguminous plants (Osdaghi et al. 2020), has had its pathogenicity linked to the presence of a linear plasmid, which contains a suite of pathogenicity-related genes (Chen et al. 2021; Evseev et al. 2022; O’Leary & Gilbertson 2020; Sparks et al. 2024; Vaghefi et al. 2021). The bacterium causes tan spot disease, also referred to as bacterial wilt or bacterial scorch (Osdaghi et al. 2020). Symptoms include leaf chlorosis and subsequent necrosis, typically initiating from leaf margins, leading to wilting, stunting and death (Diatloff & Imrie 2000; Vaghefi et al. 2021). Disease progression is most rapid at temperatures greater than 30 °C and when plants are stressed (Osdaghi et al. 2020). Seed-borne inoculum is a major source of disease, however in-field inoculum harboured on crop debris or alternative hosts may also play a role in the disease epidemiology (EPPO 2023; Gonçalves et al. 2017; Júnior et al. 2012; Osdaghi et al. 2020).

The potential for *Cff* to be hosted on a range of crops, and its risk as an emerging pathogen (EPPO 2023; Kizheva et al. 2024; Tokmakova et al. 2024), has demonstrated the need to investigate the genetic diversity among different populations, develop tools for efficient detection and explore the resources available to effectively control or reduce disease outbreaks. To support this goal, genome sequencing and population genetic studies have been conducted on *Cff* collections from Australia (Sparks et al. 2024; Vaghefi et al. 2021). These have complemented similar projects on the genetic variation within *Cff* across collections from Brazil, Iran, Russia, Spain and the United States, indicating separation into several distinct genetic clusters (Agarkova et al. 2012; Evseev et al. 2022; Gonçalves et al. 2019; McDonald & Wong 2000; Osdaghi et al. 2018). Conserved genetic regions have been identified and used for detecting the pathogen.

Species-specific PCR is an important tool for not only detecting pathogens but also quantifying pathogen load in plant tissues or field environments (Knight et al. 2022; Ophel-Keller et al. 2008; Schaad & Frederick 2002). Quantitative approaches using real-time or digital PCR platforms can support visual disease rating and provide objective measures of *in planta* pathogen populations (Knight et al. 2012; Utami et al. 2024). Applying this technique to screening plant germplasm for susceptibility to bacterial pathogens, or comparing isolate virulence, can also provide useful insights into disease traits and assist breeding programs (Garces et al. 2014; Tang et al. 2020).

Interestingly, two DNA targets for detecting *Cff*, originally reported as a restriction digestion fragment (Messenberg Guimaraés et al. 2001) or a fragment from repetitive-sequence-based PCR (rep-PCR) (Tegli et al. 2002), were based on genes now known to be in the plasmid. Studies have since reported the use of the rep-PCR sequence, predicted to be a trypsin-like serine protease gene, for *Cff* detection across several platforms (Puia et al. 2021; Tegli et al. 2020; Tegli et al. 2022). The relevance of the plasmid as a target for detection is clear, however, reports of isolates lacking the plasmid suggest the potential to under-report the presence of *Cff* in plant or environmental samples. Isolates carrying the plasmid also have the potential to vary in the number of plasmid copies (Clark et al. 2019), further complicating quantification of bacterial numbers. This issue may be alleviated by the use of PCR assays targeting conserved chromosomal regions in the *Curtobacterium* genus (Evseev et al. 2022).

The impact of *Cff* on crop production has predominantly focused on common beans (*Phaseolus vulgaris*), however its effects and survival on other crop species can have important implications for production and trade. Mung bean (*Vigna radiata* (L.) Wilczek), for example, is an increasingly popular crop due to its protein and nutrient content, short cropping duration and potential export value (Batzer et al. 2022). While mung bean production is greatest across south and southeast Asia (Pandey et al. 2018), it is emerging as a valuable crop in new regions. In Australia, mung bean is valued at an annual AUD118 million and is an expanding export commodity for the Queensland and northern New South Wales growing regions (AMA 2023; DAF 2018). Low and variable yields are a major obstacle for mung bean production in Australia, generally attributed to adverse environmental conditions and impacts of pests and diseases, such as halo blight, tan spot and powdery mildew (Batzer et al. 2022; Chauhan & Williams 2018; Kelly et al. 2021; Noble et al. 2019). Among these, tan spot has the greatest impact when the environment is hotter and dryer (Osdaghi et al. 2020), conditions that are increasingly prevalent and are already impacting agricultural production (Singh et al. 2023; Trancoso et al. 2020).

Limited information is available for *Cff* on mung bean. The earliest report of possible *Cff* in Australia occurred in 1934 on beans (*Phaseolus vulgaris*) in Victora (Adam & Pugsley 1934). Subsequently, tan spot of mung bean in Australia was first detected in 1984 across central and southern Queensland, with later reports indicating a widespread distribution (Wood & Easdown 1990). Investigations of mung bean susceptibility to tan spot, using a collection of *Cff* isolates and mung bean cultivars, demonstrated that disease expression and virulence (degree of damage) varied among the isolates and cultivars (Diatloff & Imrie 2000). With all but one cultivar reported to be susceptible, breeding efforts to introduce less susceptible germplasm continue to be a priority.

The current study aimed to explore the dynamics of the pathogenicity-related plasmid among a collection of genetically diverse Australian *Cff* isolates, including isolates not carrying the plasmid. Quantitative real-time (qPCR) and droplet digital PCR (ddPCR) assays were designed to detect and quantify the chromosome and plasmid of *Cff*, facilitating the determination of plasmid copy numbers. This knowledge was used to determine the appropriateness of the plasmid for quantitative comparisons and explore the effect of plasmid copy number variation on virulence. The disease impacts of a selection of isolates were assessed across a single susceptible mung bean cultivar. This information will be valuable for understanding the role of plasmids in pathogenicity and virulence, while also assisting with screening mung bean and related germplasm for susceptibility to *Cff*.

## 2 RESULTS

### 2.1 PCR design

Assay *Cff*_141 targeted a conserved region among the available plasmid sequences for *Cff* (Table 1; Figure S1). A BLAST search indicated the primer and probe sites targeted a conserved and specific sequence for *Cff*. The assay was re-designed from published assays to amplify a shorter fragment to ensure efficient amplification in quantitative PCR, and to include a hydrolysis probe which enabled multiplexing with other PCR targets.

**Table 1.**
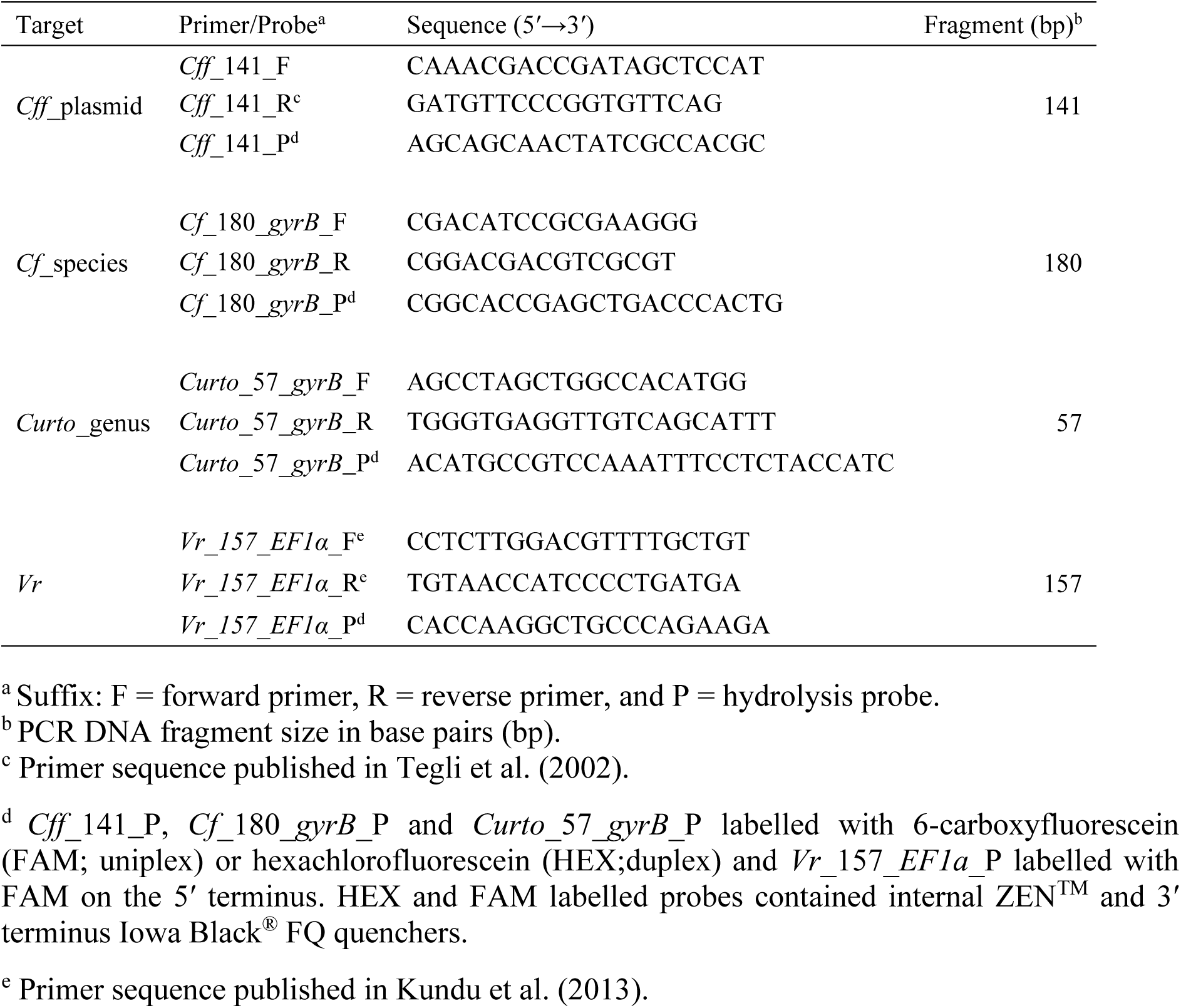
Primer/probe sets for detection and quantification of DNA sequences in the *Curtobacterium flaccumfaciens* pv. *flaccumfaciens* (*Cff*) plasmid, the gyrase B (*gyrB*) gene of *Curtobacterium flaccumfaciens* (*Cf*), the *Curtobacterium* genus (*Curto*) and the elongation factor 1-alpha (*EF1α*) gene of mung bean (*Vigna radiata*; *Vr*).

The *Cf_180_gyrB* assay target (a chromosomal species-specific region of the *Cff* gyrase B [*gyrB*] gene; Table 1) was conserved for 112 of 117 Australian *Cff* isolates, with five isolates (BRIP 70602, Cff063, Cff064, BRIP 70610 and Cff080) containing single nucleotide polymorphisms in the primer binding regions. A BLAST search indicated the primer and probe sites were partially conserved within the *C. flaccumfaciens* species. Identical primer binding site sequences were detected in other species, indicating potential cross-amplification, however the probe provided useful discrimination between *C. flaccumfaciens* and other species. Alignments across *C. flaccumfaciens* accessions suggested the assay may amplify other pathovars (Figure S2).

The *Curto*_57_*gyrB* assay target (a genus-specific region of the *Cff gyrB* gene; Table 1) was conserved for 117 Australian *Cff* isolates and most GenBank accessions for *Cff*. A BLAST search indicated the primer and probe sites were partially conserved among the *Curtobacterium* genus; however, the primer binding site sequences were identical in many other species. The probe binding site sequence was identical to predominantly the *Curtobacterium* genus. This demonstrates that the *Curto*_57_*gyrB* assay can be an alternative detection or quantification tool for *Cff* in controlled experiments, and potentially allow detection of diverse *Curtobacterium* species in mixed samples (Figure S3).

The *Vr*_157_*EF1α* assay (targeting the elongation factor 1-alpha (*EF1α*) gene of mung bean; Table 1) combined published primers for *V. mungo* (Kundu et al. 2013) with the design of a probe (Figure S4). A BLAST search indicated the primer and probe binding sites were shared amongst many species, including the single accession available for the *EF1α* gene of *V. radiata* (GenBank accession: XM_014656474). The reverse primer provided the greatest discrimination among species.

### 2.2 Plasmid copy numbers

Based on the relative quantities of ddPCR targets in the pCff119 plasmid and the gyraseB gene, there were 1.37 to 2.74 plasmid copies present in plasmid-carrying isolates (Figure 1; Table S1). Two isolates (Cff089 and BRIP 70624) contained no detectable plasmid DNA. The plasmid copy number was significantly different among the 25 *Cff* isolates (*p* <0.001; Figure 1, Table S2). The five plasmid-carrying isolates used in inoculation experiments had plasmid copy numbers of 2.38 (BRIP 70623), 2.43 (BRIP 70614), 2.45 (BRIP 70606), 2.72 (CffT13932) and 2.74 (DAR 56672). Plasmid copy numbers in DNA extracted from diseased leaves occurred across a range similar to the pure isolates (*data not shown*).

**Figure 1.**
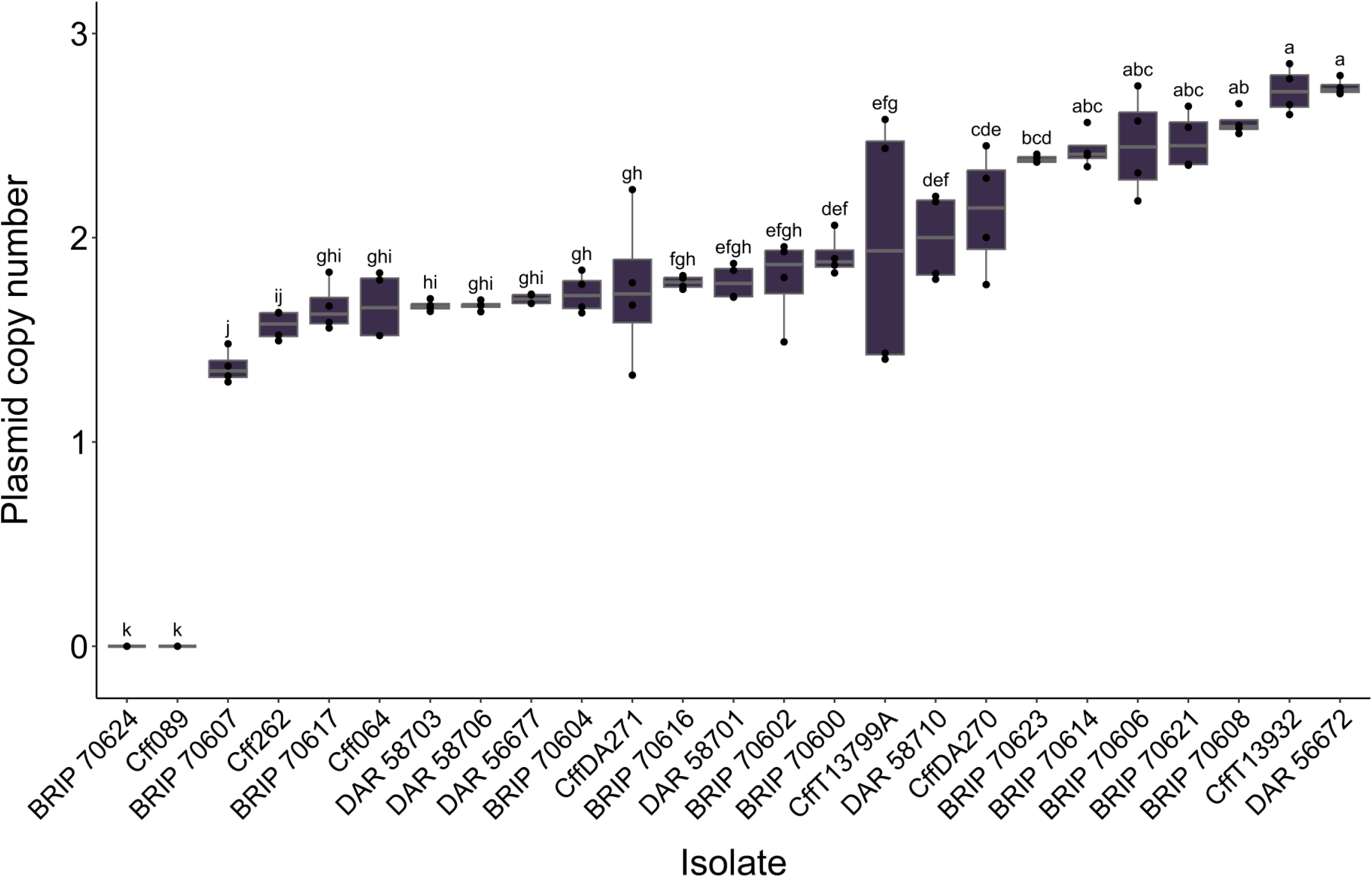
Estimated plasmid copy numbers for 25 *Curtobacterium flaccumfaciens* pv. *flaccumfaciens* isolates. The copy numbers were determined by the ratio of plasmid DNA to gyrase B DNA copies in diluted cell suspensions in droplet digital PCR. The grey centre line denotes the median value (50^th^ percentile), while the purple boxes contain the 25th to 75th percentiles of the data. The black whiskers denote the minimum and maximum values and black dots indicate the individual data points. Isolates with the same letter are not significantly different in their plasmid copy number (*n* = 4, *p* < 0.05 significance level). Mean separations were calculated on plasmid copy number values using Conover’s test.

Relative read depth comparisons from Illumina sequencing data for the pCff119 plasmid and single-copy *gyrB* and *rpoB* genes produced plasmid copy number values ranging from 1 to 5.9. Mean estimated plasmid copy numbers were similar among the ddPCR (*x̅* = 2.0, *n* = 23), *Cff*_141 to *Curto*_57_*gyrB* sequenced amplicons (*x̅* = 2.4, *n* = 113), *Cff* plasmid to *gyrB* sequenced DNA (*x̅* = 2.2, *n* = 113) and *Cff* plasmid to *rpoB* sequenced DNA (*x̅* = 2.1, *n* = 113) analyses (Table S1). The relative sequenced read depth of *gyrB* to *rpoB* ranged from 0.7 to 1.4 (*x̅* = 1.0, *n* = 119), indicating each chromosomal target is present at the same number of copies.

### 2.3 Quantitative PCR characteristics

The *Cf_180_gyrB* assay was able to detect *Cff* DNA from 0.25 pg to 25 ng and the *Vr*_157_*EF1α* assay was able to detect mung bean DNA from 25 pg to 25 ng. A standard curve was calculated for the *Cf_180_gyrB* assay (*R^2^* = 0.99, *y* = −3.6*x* + 15.7, *E* = 89.9) and the *Vr*_157_*EF1α* assay (*R^2^* = 0.99, y = −3.2*x* + 28.1, *E* = 105.0). Based on these results a cut-off point, or limit of quantification, was set at 31.6 cycles for *Cff* DNA. The respective mean values (*n* = 8) and coefficient of variation (CV) of a mixed DNA sample assessed for intra-assay variability were 1.8 ng *Cff* DNA (CV = 10.2%) for *Cf_*180*_gyrB* and 31.0 ng mung bean DNA (CV = 21.8%) for *Vr*_157_*EF1α*. The respective mean values (*n* = 4) and CV of the mixed DNA sample assessed for inter-assay variability were 2.0 ng *Cff* DNA (CV = 11.1%) for *Cf_*180*_gyrB* and 29.2 ng mung bean DNA (CV = 13.0%) for *Vr*_157_*EF1α*.

### 2.4 Tan spot disease symptoms

Mung bean plants were inoculated with seven treatments, including five plasmid-carrying isolates, one plasmid-free isolate and a water control (Table 2). Initial disease symptoms were observed on the trifoliate leaves three days after inoculation, with the timing of symptom expression dependent on isolate (Figure 2). Visual disease rating at 15 days after inoculation was based on summed values for three trifoliate leaves, with potential scores ranging from 0 to 300%. Significant variation in the expression of visual disease symptoms occurred among the *Cff* isolates (*p* <0.001) (Figure 3A, Table S3). For the plasmid-carrying isolates, the mean visual symptoms ranged from 266.1% for the DAR 56672 treatment to 127.5% for the BRIP 70623 treatment. No symptoms were observed on the mung bean plants inoculated with the plasmid-free isolate Cff089 or the water treatment.

**Figure 2.**
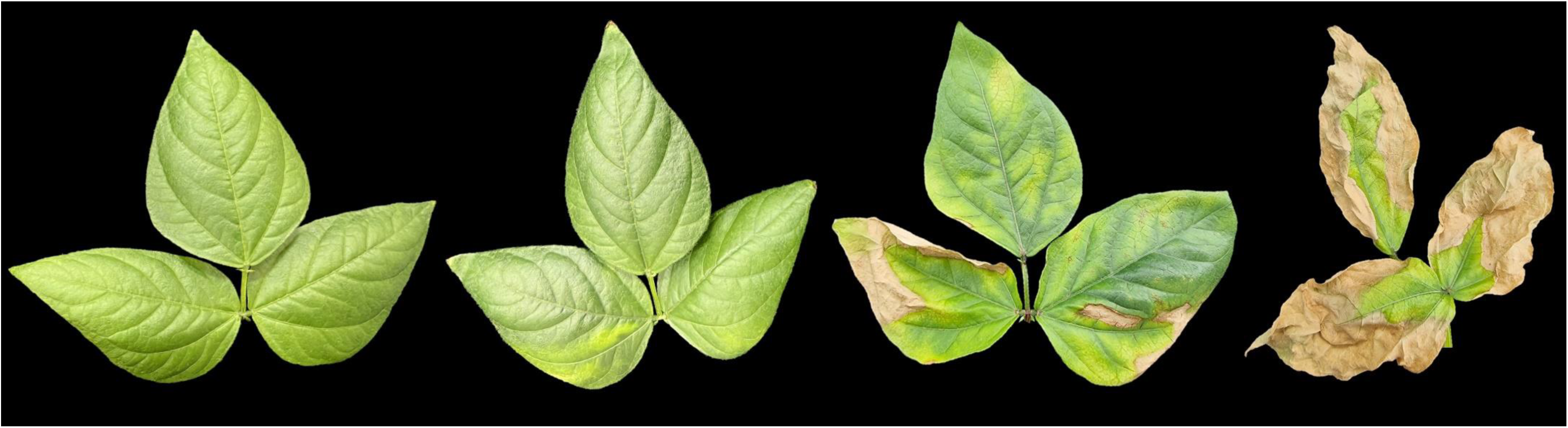
Example of tan spot symptoms on the first trifoliate leaf of mung bean cultivar Opal-AU. Leaf responses range from no symptoms to 100% symptomatic leaf tissues (left to right). Visual rating accounted for both chlorosis and necrosis across the leaf area. Images were edited in PowerPoint, Microsoft 365.

**Figure 3.**
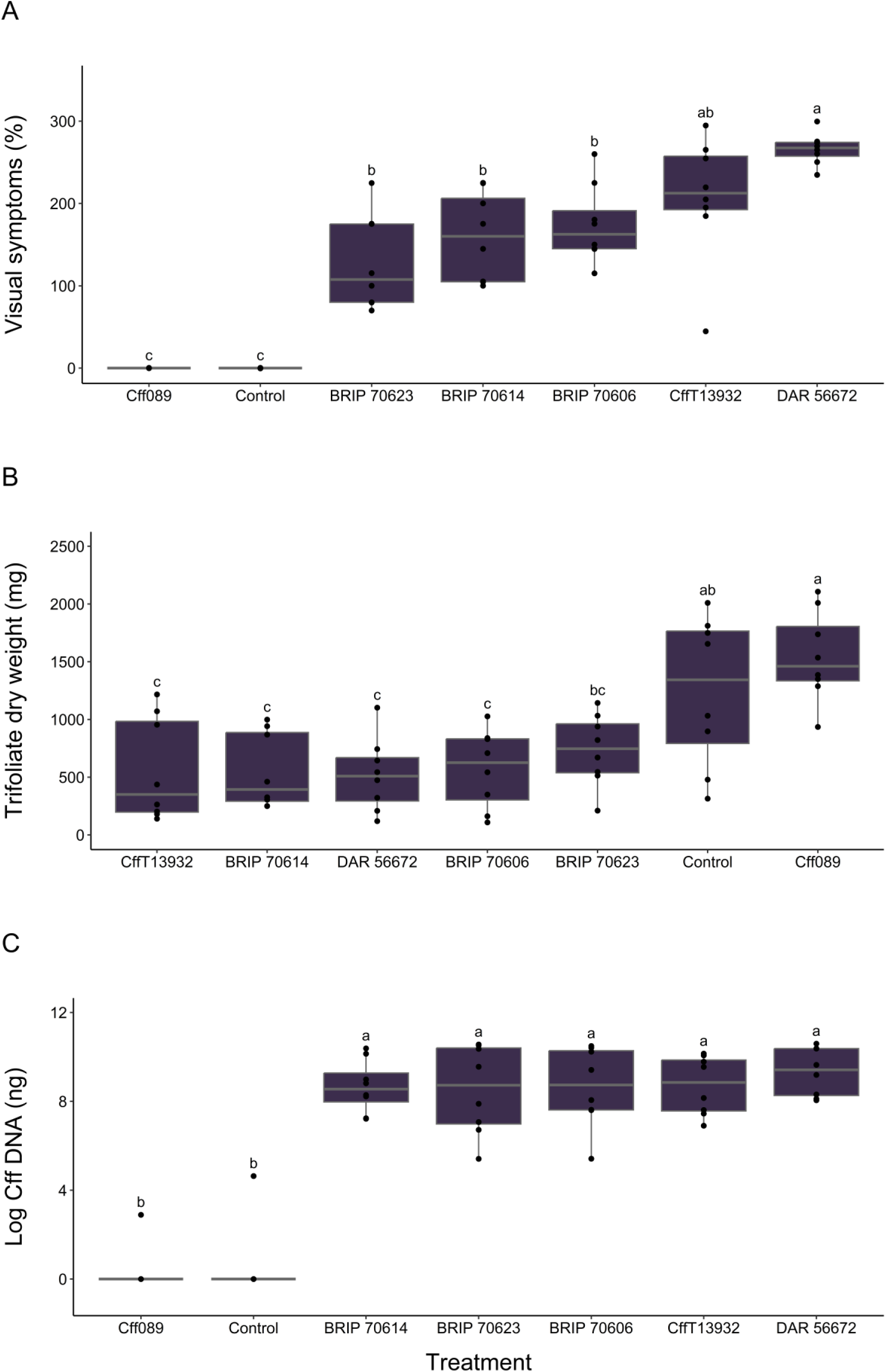
Disease responses caused by six *Curtobacterium flaccumfaciens* pv. *flaccumfaciens* (*Cff*) isolates and a water control treatment at 15 days after inoculation of mung bean plants (cv. Opal-AU, *n* = 6). Responses include visual leaf disease symptoms (A), trifoliate leaf dry weights (B) and *Cff* DNA (log[x+1]) in total leaf tissue samples (C). Different letters represent significant differences among the treatments at *α* < 0.05. Mean separations were calculated using Dunn’s test.

**Table 2.**
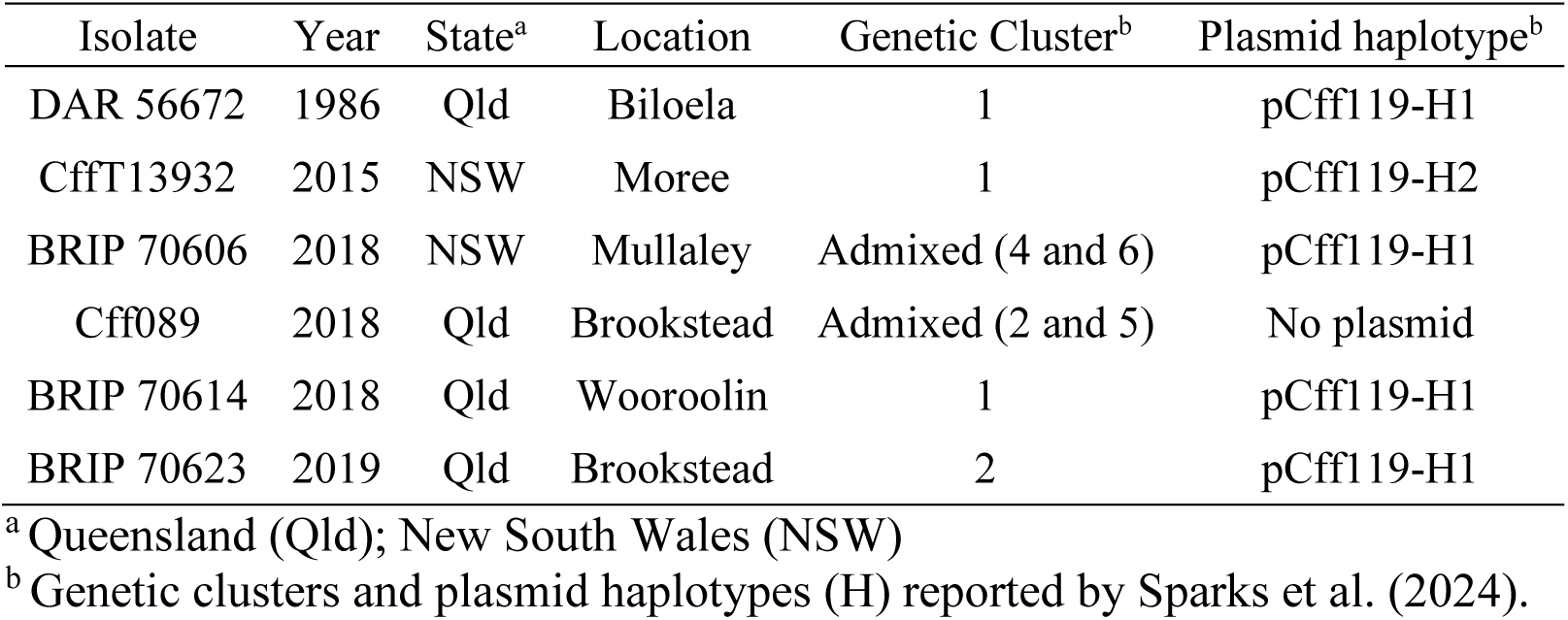
Metadata for six *Curtobacterium flaccumfaciens* pv. *flaccumfaciens* isolates selected for mung bean inoculation experiments. Each isolate was originally isolated from mung bean.

### 2.5 Trifoliate leaf dry weight

Inoculation with each of the five plasmid-carrying isolates significantly reduced the trifoliate dry weight compared to the control plants (*p* <0.002). Visible stunting in the height of these plants was also observed. Among these plants, the mean trifoliate leaf dry weights ranged from 571 mg for the BRIP 70606 treatment to 520 mg for the DAR 56672 treatment, with no significant difference among the four treatments. Treatment BRIP 70623 (735 mg) was not significantly different from the control (1244 mg) or the other plasmid-carrying isolate treatments. In contrast, the trifoliate leaf dry weights of plants inoculated with the plasmid-free isolate Cff089 (1544 mg) were 19.7% greater than the control plants. However, this difference was not significant at *α* = 0.05 (Figure 3B, Table S4).

### 2.6 Curtobacterium flaccumfaciens pv. flaccumfaciens DNA quantities

Significant variation in the *Cff* DNA quantities was reported among the seven treatments (*p* <0.001) (Figure 3C, Table S5). This was predominantly due to the Cff089 and control treatments, for which only one sample contained a detectable quantity of *Cff* DNA (17 ng and 102 ng, respectively). The remaining Cff089 samples had quantification cycle (Cq) values outside the range of quantification (31.87 to 36.69). For the remaining control samples, two had no detection, with the other five samples exhibiting Cq values outside the range of quantification (31.71 to 38.02). The control sample with detectable *Cff* DNA was associated with a plant observed to have mild visible symptoms, not initially ascribed to tan spot disease but likely due to inoculum crossover in the randomised blocks. No significant difference was observed for the *Cff* DNA quantity among the plasmid-carrying isolate treatments, with the mean values ranging from 17,509 ng for the DAR 56672 treatment to 10,376 ng for the BRIP 70614 treatment.

### 2.7 Correlation of disease traits

A significant positive correlation occurred between visual disease symptoms and *Cff* DNA quantity (*ρ* = 0.66, *p* < 0.001). A significant negative correlation occurred between trifoliate leaf dry weights and visual disease symptoms (*ρ* = −0.61, *p* < 0.001) and *Cff* DNA quantity (*ρ* = −0.41, *p* = 0.002), respectively.

## 3 DISCUSSION

Tools for detecting and quantifying genetic characteristics of pathogens are important for disease assessment and understanding pathogen dynamics. This study reports the development of PCR-based assays for the quantification of chromosomal- and plasmid-associated regions of the *Cff* genome. These assays were used to provide the first demonstration of variation in the plasmid copy number among *Cff* isolates. The chromosome-based PCR assay also allowed detection and quantification of a single-copy chromosomal region in all *Cff* isolates, regardless of plasmid presence or absence. These tools demonstrated variation in disease traits on mung bean among a selection of six *Cff* isolates. This approach and information will be beneficial for assessing and managing disease on crops affected by *Cff*.

The presence of a plasmid containing pathogenicity-related genes has been linked to pathogenicity in *Cff* (Chen et al. 2021; Evseev et al. 2022; O’Leary & Gilbertson 2020; Osdaghi et al. 2020; Sparks et al. 2024; Vaghefi et al. 2021). However, non-pathogenic isolates without the plasmid have also been isolated from diseased plant tissues (Vaghefi et al. 2021). Bacteria can exchange genetic material, which may include the loss or gain of a plasmid and subsequent alterations of fitness in different environments (Vivian et al. 2001). The existence of *Cff* isolates without the plasmid demonstrated a need for alternative PCR detection methods beyond those which have been reported for *Cff* detection based on a region found in the plasmid (Messenberg Guimaraés et al. 2001; Tegli et al. 2002). This led to the development of PCR assays based on the *gyrB* gene, which successfully amplified for the majority of *Cff* isolates, whilst also theoretically discriminating against other bacteria. Due to DNA sequence variability, a single assay capable of detecting all *Cff* isolates, whilst also discriminating against other related bacteria, could not be generated.

Plasmid copy number was not fixed among the isolates. This was determined by directly testing bacterial suspensions in ddPCR, thus avoiding bias due to DNA extraction procedures (Plotka et al. 2017). The findings were confirmed with read depth analyses of genome sequence data. This provided further justification for the development of alternative quantitative PCR assays. Single-copy chromosomal genes, such as *gyrB*, are preferred targets for DNA quantification via PCR, however the plasmid and chromosome of *Cff* are physically independent and the observation suggested they did not occur in a 1:1 ratio. This has implications for reporting DNA quantities and estimating bacterial load. For example, a 2:1 plasmid to chromosome ratio may result in twice as much bacterial load being reported when using a plasmid-related PCR assay. If the plasmid copy number varies among isolates, then relevant comparisons would be impossible.

The multicopy nature of the plasmid lends itself to a greater chance of detection, providing a benefit for presence/absence screening. However, comparative PCR quantification based on a plasmid target should be avoided. The epidemiological role of variation in plasmid copy number among *Cff* isolates is not yet established but has potential implications for disease expression. Interestingly, isolate DAR 56672 exhibited the greatest plasmid copy number, whilst also being associated with the greatest values for visual symptom expression, bacterial biomass production and plant biomass reduction. It is possible that greater plasmid copy numbers may influence the strength of plasmid-borne characteristics (Clark et al. 2019; Friehs 2004; Schierstaedt et al. 2019) or increase the rate of plasmid transmission (Dimitriu et al. 2021; Williams & Thomas 1992), although plasmids may also confer a fitness cost (San Millan & MacLean 2017; Williams & Thomas 1992). Virulence in the human pathogen *Yersinia pestis* has been linked to up-regulated plasmid copy numbers during infection, however plasmid copy numbers in plant pathogens have received limited attention (Coplin 1989; Schierstaedt et al. 2019; Vivian et al. 2001). Plasmid copy numbers may also be linked to the process of bacterial replication, with greater plasmid copy numbers increasing the chance of transferal to daughter cells and the inheritance of pathogenicity (Jahn et al. 2015). Retention of plasmids occurring at lower numbers may also involve active partitioning to decrease the chance of a plasmid-free cell developing (Williams & Thomas 1992).

Plasmid-free isolates and their lack of pathogenicity have been documented in previous reports (Sparks et al. 2024). The pathogenicity function of the plasmid was further evidenced in the current study by the non-pathogenic plasmid-free isolate Cff089. Not only did this isolate not induce visual disease symptoms or reduce trifoliate dry weights, it also did not proliferate in the mung bean tissues asymptomatically. This leads to questions regarding the role of such isolates in the environment and whether they may pose a disease risk if the plasmid is acquired, while also providing opportunities to study the influence of the plasmid on bacterial growth characteristics. It is possible that the population from which isolate Cff089 was derived had the plasmid and was pathogenic but the plasmid was lost during cell division, either in the plant tissue or during successive culturing following the concomitant relaxation of selection pressures for its retention (Smith & Bidochka 1998). The complex situation of *Cff* isolates ranging from no plasmid to variation in plasmid copy numbers, along with variation in pathogenicity and virulence, suggests a need to elucidate the role of the genes in the plasmid.

Disease reactions to *Cff* have traditionally been assessed based on visual inspections (Diatloff & Imrie 2000; Harveson et al. 2015; Wood & Easdown 1990). While these can be relatively rapid and cheap to perform, they are subjective and limited by the ability to visually characterise precise disease symptoms, which can lead to variation among experiments. Image-based analysis has also been applied to *Cff* disease symptoms (Vaghefi 2022), improving the precision of ratings but increasing the time to assess symptoms. In contrast, PCR-based methods of quantification provide objective measures of pathogen numbers and can be standardised by using a consistent tissue weight for DNA extraction or quantifying a host DNA target (Knight et al. 2012; Utami et al. 2024). Plant tissue dry weights also provide an objective assessment of the impact of a pathogen on plant growth, offering a further perspective on disease impacts. While useful to apply, both PCR-based and plant tissue dry weight methods require more resources than traditional visual rating. Strategic application of alternative disease assessment methods can support visual assessments and offer further insights into plant-pathogen interactions.

The assessment of tan spot disease traits on mung bean cultivar Opal-AU suggested variation among the plasmid-carrying isolates for visual symptoms. In contrast, *Cff* DNA quantities and trifoliate dry weights provided no significant distinction, but a trend was observed where isolate DAR 56672 caused the most severe disease responses. Sparks et al. (2024) similarly reported significant differences in disease severity among 11 *Cff* isolates on the moderately resistant mung bean cultivar Jade-AU. A previous study by Diatloff and Imrie (2000) identified mild and severe disease causing isolates of *Cff*, however, upon inoculation across a range of mung bean cultivars, it was reported that their relative virulence varied. This isolate by cultivar interaction suggests that the use of a single susceptible cultivar in the current study may have limited the potential to discriminate among isolates for their disease impacts.

Assessment of visual symptoms, *Cff* DNA quantities and trifoliate dry weights facilitated consideration of a range of disease impacts. The consistent reduction of trifoliate dry weights suggests a general plant response to each isolate, with a significant reduction in biomass suggested to correlate to reduced grain yields (Geetika et al. 2022). In contrast, a range of visual disease symptoms was observed, indicating variation in foliar responses to each isolate. The trifoliate dry weight and visual observations may be related to the stem inoculation method, which delivered an equal quantity of bacteria to each plant to incite a growth response, and the subsequent ability of each isolate to move within the plant tissue.

The assessment of *Cff* DNA quantities provides another perspective of the disease process, with a focus primarily on the ability of the bacteria to move to, colonise and reproduce in the trifoliate tissues. The *Cff* DNA quantities suggested a range among the isolates, however no statistically significant differences occurred. While PCR-based biomass estimates may exhibit variation due to the rate of disease expression and pathogen reproduction, there is a potential impact of *Cff* asymptomatically colonising mung bean tissues (Diatloff & Imrie 2000). Any asymptomatic colonisation may have reduced the ability to discriminate among the isolates based on DNA quantities. Assessing three disease traits provides a more comprehensive perspective of the interactions between the host and the pathogen, while also suggesting that objective continuous variables, such as *Cff* DNA quantities and trifoliate dry weights, provide both challenges for comparative assessments and precise interpretations of biological variation.

The information from this study begins to unravel the plasmid dynamics in *Cff* and indicates a need to further investigate the regulation of plasmid replication, inheritance and transmission, along with expression of genes in the plasmid, and the relationship between plasmid copy number and virulence. The alternative PCR tools for *Cff* detection and quantification may be used to explore disease across different crops affected by *Cff* and investigate inoculum sources. Furthermore, the classification of virulence among a group of genetically diverse isolates will be used to support mung bean breeding efforts to screen for reduced susceptibility to *Cff*.

## 4 EXPERIMENTAL PROCEDURES

### 4.1 PCR assay design

Four probe-based PCR assays for real-time quantitative PCR (qPCR) or ddPCR platforms were developed to amplify a plasmid-specific region in *Cff* (assay *Cff*_141), a species-specific region of the *Cff* gyrase B (*gyrB*) gene (assay *Cf*_180_*gyrB*), a genus-specific region of the *Cff gyrB* gene (assay *Curto*_57_*gyrB*) and the elongation factor 1-alpha (*EF1α*) gene of mung bean (assay *Vr_157_EF1α*) (Table 1). Alignments and primer and probe design were performed against reference sequences as described by Knight et al. (2022).

The plasmid-specific assay was a modified design based on a region targeted by Tegli et al. (2002). Modification included reducing the amplicon size and designing a probe. The targeted sequence was aligned across five published plasmid sequences (GenBank accessions CP080396, CP041260, CP045288, CP071884 and CP074440).

Due to reports of *Cff* isolates without a plasmid (Sparks et al. 2024; Vaghefi et al. 2021) and variation in plasmid copy numbers among isolates, assays were designed based on the single-copy phylogenetically-informative chromosomal gene *gyrB*. Sequences of 117 *Cff* isolates from Australia (Sparks et al. 2024), *Cff* type strain DSM 20129 (GenBank Accession: CP080395) and closely related species and pathovars were used for primer and probe alignment.

Amplification of mung bean DNA used primers described by Kundu et al. (2013) and a probe designed against the *EF1α* sequence of *V. radiata* (GenBank accession XM_014656474). Mung bean DNA detection acted as an internal positive control in PCR tests of plant DNA templates.

### 4.2 Plasmid copy number assessment

The copy number of the linear plasmid (pCff119) (Vaghefi et al. 2021) was assessed for 25 *Cff* isolates (Table S1) using a duplex PCR reaction of the *Cff*_141 (plasmid) and *Curto*_57_*gyrB* (chromosome) assays in a QX200 ddPCR platform (Bio-Rad Laboratories, California, USA), following the manufacturer’s instructions. Each assay was initially optimised in uniplex across a temperature gradient (55.5 to 65.5 °C) before combining as a duplex. Each 22 μL ddPCR reaction contained 11 µL of 2× ddPCR Supermix for Probes (no dUTP) (Bio-Rad Laboratories), 0.25 µM of each forward and reverse primer (Integrated DNA Technologies, Iowa, USA), 0.15 µM of each probe (Integrated DNA Technologies) and 5 µL of template. Thermal cycling conditions consisted of 95 °C for 10 min followed by 50 cycles of 94 °C for 30 s and 58 °C for 60 s, with a final denaturation at 98 °C for 10 min, followed by a hold step at 4 °C. Copy numbers for each target were calculated in the QX Manager Standard Edition software v. 1.2 (Bio-Rad Laboratories).

Control templates included *Cff* DNA (0.5 ng) as a positive control and sterile water as a no-template control (NTC). The experimental samples were *Cff* cell suspensions of 1.0 OD at 600 nm, grown as described for inoculum production below, and adjusted with a 10^5^-fold dilution. Cell suspensions were stored overnight at −20 °C before testing. Representative DNA samples from the plant inoculation experiments described below were also assessed. Each template was assessed in duplicate.

A BLAST search of the *Cff* genome (taxid:138532) confirmed the presence of only one sequence target for each PCR assay in the plasmid and chromosome, respectively. Therefore, for each ddPCR reaction, the relative copy number of the *Cff*_141 (plasmid) PCR target was calculated by dividing the ddPCR copy numbers of the *Cff*_141 assay by the ddPCR copy numbers of the Curto_57_*gyrB* assay. The four values (two technical replicates of two biological replicates) were used to produce a boxplot with the package *ggplot2* version 3.4.2 in RStudio version 2023.06.0 (R Core Team 2017).

Plasmid copy numbers were also assessed based on relative read depths to the single-copy genes *gyrB* and *rpoB* in Illumina paired-end sequences of 119 isolates of *Cff* from Australia (Sparks et al. 2024). Sequence reads for each isolate were independently aligned to sequences of a complete plasmid (pCFF113; NCBI Accession: CP041260.1), a partial *gyrB* gene (NCBI Accession: MW732638.1) and a *rpoB* gene (NCBI Accession: KT935421.1) using the short-read aligner Burrows-Wheeler Aligner (bwa-mem) version 0.7.15 with default settings (Li & Durbin 2009). SAMtools version 1.11 (Li et al. 2009) was used to sort and convert data from Sequence Alignment Map into Binary Alignment Map formats. The “depth” function in SAMtools (with the -a option for plasmid/gene sequences, or the -a and -r options for PCR amplicon regions) was used to calculate read depth (the number of sequence reads covering each specific base pair position in the reference sequence) across the plasmid/gene sequences and the *Cff*_141 and *Curto*_57_*gyrB* amplicon sequences. Finally, the plasmid size and total read depth were calculated, and the “statistics” function in Python was used to determine the mean read depth. The relative copy numbers of the plasmid were calculated by dividing the read depth of the plasmid sequence by the read depths of the *gyrB* or *rpoB* gene regions.

### 4.3 Quantitative real-time PCR conditions

Quantitative real-time PCR was conducted on a CFX Opus 96 or CFX384 platform (Bio-Rad Laboratories). Assays were initially assessed in uniplex, across a gradient of annealing temperatures (48 to 66 °C) and with target and non-target DNA (0.5 ng/µL) (Table S6). Duplex assays of *Cff*_141, *Cf*_180_*gyrB* or *Curto*_57_*gyrB* and *Vr_157_EF1α* were also assessed. In uniplex reactions all probes were labelled with a 5′ terminus 6-carboxyfluorescein (FAM) fluorophore, while in duplex reactions the *Vr_157_EF1α* probe remained labelled with a FAM fluorophore and each bacterial probe was labelled with a 5′ terminus hexachlorofluorescein (HEX) fluorophore. All probes contained internal ZEN^TM^ and 3′ terminus Iowa Black^®^ RQ quenchers. Each 20 µL reaction consisted of 10 µL of ImmoMix (Bioline, London, UK), 0.25 μM of each forward and reverse primer (Integrated DNA Technologies), 0.15 μM of probe (Integrated DNA Technologies) and 5 μL of template DNA. Optimal thermal cycling conditions consisted of 10 min at 95 °C, followed by 40 cycles of 95 °C for 15 s and 62 °C for 60 s.

The duplex of *Cf*_180_*gyrB* and *Vr_157_EF1α* was used to test the inoculated mung bean DNA samples. For quantification, a ten-fold dilution series of *Cff* DNA ranging from 25 ng to 0.025 pg per reaction (final volume) was assessed in triplicate. A standard curve was generated using the CFX Maestro Software v. 1.1 (Bio-Rad Laboratories) by plotting the logarithm of *Cff* DNA concentrations against the quantification cycle (Cq). Pure *Cff* and mung bean DNA templates were included as positive and negative controls, with sterile water included as a NTC. A mixed DNA template containing both *Cff* and mung bean DNA (extracted from Opal-AU tissue inoculated with isolate BRIP 70623) was also included in each run. Each sample template was assessed in duplicate.

Mean *Cff* DNA values were transformed to indicate the total DNA quantity in the entire dry trifoliate sample for each plant (Knight et al. 2012). Briefly, for each inoculated mung bean DNA template, the mean DNA quantity value was multiplied by 10 (to account for the ten-fold dilution of the original DNA sample), divided by 5 to get the ng/µL value, then multiplied by 100 (microlitre volume of eluted DNA) to determine the total quantity of *Cff* DNA extracted from the sub-sample. The total *Cff* DNA quantity was divided by the sub-sample weight to determine the *Cff* DNA quantity per milligram, then multiplied by the total trifoliate leaf dry weight to determine the *Cff* DNA quantity in the combined trifoliate leaf sample.

For the *Cf*_180_*gyrB* and *Vr*_157_*EF1α* duplex reaction, intra- and inter-assay variation were reported as the mean DNA quantity and coefficient of variation (CV) of replicate samples of a mixed *Cff* and mung bean DNA template.

### 4.4 Planting and glasshouse conditions

Two replicated glasshouse trials were conducted at the University of Southern Queensland, Australia in 2022 and 2023. Mung bean cultivar Opal-AU, considered susceptible to tan spot (Douglas 2021), was grown for pathogenicity experiments. Two seeds were planted into 1.5 L pots (14 cm diameter) containing steam-sterilised (75 °C for 45 min) potting mix (Scotts Osmocote Premium Plus Superior, Evergreen Garden Care Australia, New South Wales, Australia). Pots were placed on metal trays and kept under natural light conditions, with day/night temperatures of 27 to 33 °C and 20 to 25 °C, respectively. Pots were manually watered at the soil surface prior to inoculation, with watering post-inoculation via submersion of the pot bases in approximately 1 cm of water twice a week. After germination, each pot was thinned to one plant. At seven days after planting pots were fertilised with Seasol ‘Complete Garden Health Treatment’ (Seasol International Pty Ltd, Victoria, Australia) at 0.7 % (v/v) and at two weeks with Season PowerFeed (Seasol International Pty Ltd) at 1.1 % (v/v) in rainwater (Vaghefi 2022).

### 4.5 Inoculum production, inoculation and harvest

Six *Cff* isolates, collected in a previous study (Sparks et al. 2024), were selected for pathogenicity and virulence assessment (Table 2). Selection was informed by genetic clustering (Sparks et al. 2024) and temporal and geographic origin. Isolates were grown on Nutrient Agar (Amyl Media, Victoria, Australia) for three days and a single colony was transferred to 10 mL of Nutrient Broth (NB, Amyl Media, Victoria, Australia) in a 50 mL tube before placing in a shaker-incubator at 180 rpm and 27 °C for 48 h. On the day of inoculation, each culture was centrifuged for 10 min at 11,200 g, the supernatant discarded, and the bacterial pellet re-suspended in 8 mL of sterile water. Each cell suspension was diluted to an optical density (OD) of 1.0 at 600 nm (DS-11FX spectrophotometer, DeNovix Inc, Delaware, USA), which is equivalent to 1.0 × 10^8^ CFU mL^−1^.

The inoculation experiment included seven treatments, consisting of the six isolates and a sterile water control. Inoculations were conducted after the emergence of the first trifoliate leaf, approximately 15 days after planting. For each treatment, 10 μL of cell suspension or water was placed on the stem between the unifoliates and the base of the first trifoliate leaf petiole. An acupuncture needle (0.3 mm diameter, Dongbang Medical Co, Ltd, South Korea) was used to pierce through the stem and inoculum droplet three times. The experiment was performed twice, and each was arranged in a randomised complete block design with four replicates for each of the seven treatments.

At 15 days after inoculation, the first three trifoliate leaves were visually rated for symptom expression. Visual rating used a 0 to 100% scale (5% increments) to measure the chlorosis and necrosis visible on each leaf, where 0 indicates no symptom development and 100 indicates expression of chlorosis and/or tan-brown necrosis across the entire leaf area (Figure 1). The sum of the three trifoliate visual ratings for each plant was used for subsequent analyses. The first three trifoliate leaves of each plant were harvested and placed in a paper bag. Tissues were dried for three days at 55 °C and weighed.

### 4.6 Tissue preparation and DNA extraction

Dried leaf tissues were placed into 15 mL tubes with two steel balls (4.5 mm diameter) and ground at 6.5 m/s for 60 s in a FastPrep-24 instrument (MP Biomedicals, California, USA). A sub-sample of 15 to 20 mg was taken and DNA extracted using the DNeasy Plant Mini Kit (Qiagen, Hilden, Germany) according to the manufacturer’s instructions. To reduce the possible impact of inhibitors, each sample was diluted ten-fold in sterile water.

Pure *Cff* DNA was extracted from cells grown in NB (as described above) using the GenElute Bacterial Genomic DNA Kit (Merck Pty Ltd, Darmstadt, Germany) according to the manufacturer’s instructions. Pure mung bean DNA was extracted from one-week old leaf tissue using the DNeasy Plant Pro Kit (Qiagen). DNA of non-target organisms (Table S6), used for PCR optimisation, was extracted from pure tissues using the DNeasy Plant Mini Kit (Qiagen). DNA characteristics were measured using a DS-11FX spectrophotometer (DeNovix Inc) and quantified using a Qubit 3.0 Fluorometer (Thermo Fisher Scientific, Massachusetts, USA).

### 4.7 Statistics

All statistical analyses were performed in RStudio version 2023.06.0 (R Core Team 2017; Wickham 2016). To determine the appropriate statistical tests, the normality and homogeneity of variances of the data was assessed. The Shapiro-Wilk test (*α* = 0.05) (“shapiro.test” function in package *stats*) was used to test for normality. Homogeneity of variances (*α* = 0.05) across treatments was evaluated using Levene’s test (“leveneTest” function in package *car*). Based on these results, non-parametric methods were appropriate for analysis of plasmid copy number, visual symptom, *Cff* DNA quantity and trifoliate leaf dry weight data. Due to the wide range of *Cff* DNA quantity values, a log(x+1) transformation was applied. Independent replicated experiments were compared using the Wilcoxon rank-sum test (“wilcox.test” function in package *stats*). To compare measurements for multiple isolates or treatments, the Kruskal-Wallis test was conducted (“kruskal.test” function in package *stats*). This non-parametric method tests whether samples originate from the same distribution. Following a significant Kruskal-Wallis test (*α* = 0.05), Dunn’s test was performed for the visual symptom, *Cff* DNA quantity and trifoliate leaf dry weight data to conduct pairwise comparisons (“dunn.test” function in package *dunn.test*). For the plasmid copy number data, Conover’s test was used to generate pairwise comparisons (“conover.test” function in package *conover.test*). These tests identify specific group differences and adjust for multiple comparisons using the Benjamini-Hochberg method (Benjamini & Hochberg 1995). Significant differences identified by Dunn’s test or Conover’s test were summarised using a compact letter display (“cldList” function in package *multcompView*), which assigns letters to groups based on their statistical significance. Boxplots were generated to visualise the distribution of each data set (function “geom_boxplot” in package *ggplot2*). Correlations between visual symptoms, *Cff* DNA quantity and trifoliate leaf dry weight were assessed using Spearman’s rank correlation coefficient (“cor.test” function in package *stats*).

## ACKNOWLEDGEMENTS

This research was funded by the Broadacre Cropping Initiative (Project 36), a joint investment of the University of Southern Queensland and the Queensland Government Department of Primary Industries. The authors thank C. A. Douglas for discussions during project development and supplying Opal-AU seed, and K. Cowan (New South Wales Plant Pathology and Mycology Herbarium), Y. P. Tan (Queensland Department of Primary Industries Plant Pathology Herbarium), D. L. Adorada and E. E. Adorada (University of Southern Queensland) for supplying and creating bacterial and fungal isolates.

## DATA AVAILABILITY STATEMENT

The data that support the findings of this study are available from the corresponding author upon reasonable request.

