## Supplementary material for "Plasmid copy number variation impacts pathogenicity and quantification of *Curtobacterium flaccumfaciens* pv. *flaccumfaciens* infecting mung bean": Table S

### SUPPLEMENTARY TABLES AND FIGURES

**Table S1.** *Curtobacterium flaccumfaciens* pv. *flaccumfaciens* (*Cff*) plasmid pCff119 copy numbers estimated using droplet digital PCR (ddPCR) ($n$ = 25 isolates) and relative sequence read depths across PCR amplicons and entire plasmid and gene (*gyrB* and *rpoB*) regions ($n$ = 119). ddPCR estimated plasmid copy numbers by dividing the *Cff*_141 by *Curto*_57_*gyrB* copies across four replicates, with the mean value and standard error (SE) calculated. For Illumina sequencing data either the target amplicon for the *Cff*_141 and *Curto*_57_*gyrB* assays or the entire plasmid and gene regions for *gyrB* and *rpoB* were assessed. Each were assessed for total read depth across the sequence and the mean read depth per base across the same sequence. The relative read depth, or relative copy number, was calculated using the mean read depth for the plasmid compared to both *gyrB* and *rpoB*. The *gyrB* and *rpoB* regions were also compared for relative copy number. Isolates are ordered by the relative read depth of *Cff* plasmid to *gyrB*. Table code ([a]) indicates isolates used in mung bean (cv. Opal-AU) inoculation experiments, (-) indicates isolates were not tested, (na) indicates the calculation was not applicable due to no plasmid being detected.

| Isolate | Year | State | Location | Host | Estimated plasmid copies (ddPCR) | SE (ddPCR) | Total read depth across plasmid *Cff*_141 PCR amplicon (142bp) | Total read depth across *Curto*_57_*gyrB* PCR amplicon (57bp) | Mean read depth across plasmid *Cff*_141 PCR amplicon | Mean read depth across *Curto*_57_*gyrB* PCR amplicon | Relative read depth of plasmid to *gyrB* amplicon | Length of *Cff* plasmid (bp) | Total mapped read depth across *Cff* plasmid | Mean mapped read depth across *Cff* plasmid | Length of partial *gyrB* (bp) | Total mapped read depth across *gyrB* gene | Mean read depth across *gyrB* gene | Length of *rpoB* (bp) | Total mapped read depth across *rpoB* gene | Mean read depth across *rpoB* gene | Relative read depth of *Cff* plasmid to *gyrB* | Relative read depth of *Cff* plasmid to *rpoB* | Relative read depth of *gyrB* to *rpoB* |
|---|---|---|---|---|---|---|---|---|---|---|---|---|---|---|---|---|---|---|---|---|---|---|---|
| Cff019; BRIP70603 | 2018 | NSW | Mullaley | *Vigna radiata* | - | - | 14700 | 6427 | 103.5 | 112.8 | 0.9 | 105088 | 11391630 | 108.4 | 865 | 95335 | 110.2 | 328 | 36661 | 111.8 | 1.0 | 1.0 | 1.0 |
| Cff021; BRIP 70604 | 2018 | NSW | Mullaley | *V. radiata* | 1.7 | 0.049 | 25746 | 7550 | 181.3 | 132.5 | 1.4 | 105088 | 15067724 | 143.4 | 865 | 111088 | 128.4 | 328 | 39616 | 120.8 | 1.1 | 1.2 | 1.1 |
| CffT13802B; BRIP 70619 | 2014 | Qld | Warwick | *V. radiata* | - | - | 18533 | 6314 | 130.5 | 110.8 | 1.2 | 113440 | 14168612 | 124.9 | 865 | 93104 | 107.6 | 328 | 38133 | 116.3 | 1.2 | 1.1 | 0.9 |
| CffT13802A | 2014 | Qld | Warwick | *V. radiata* | - | - | 14697 | 6221 | 103.5 | 109.1 | 0.9 | 113438 | 13492117 | 118.9 | 865 | 85955 | 99.4 | 328 | 31494 | 96.0 | 1.2 | 1.2 | 1.0 |
| CffT14069; BRIP 70620 | 2015 | Qld | Ayr | *V. radiata* | - | - | 41365 | 12938 | 291.3 | 227.0 | 1.3 | 113440 | 32868122 | 289.7 | 865 | 202701 | 234.3 | 328 | 75713 | 230.8 | 1.2 | 1.3 | 1.0 |
| Cff130; BRIP 70614[a] | 2018 | Qld | Wooroolin | *V. radiata* | 2.4 | 0.046 | 61377 | 16082 | 432.2 | 282.1 | 1.5 | 113436 | 19531355 | 172.2 | 865 | 117788 | 136.2 | 328 | 42255 | 128.8 | 1.3 | 1.3 | 1.1 |
| Cff103 | 2018 | Qld | Brookstead | *V. radiata* | - | - | 22882 | 6248 | 161.1 | 109.6 | 1.5 | 113435 | 15162303 | 133.7 | 865 | 89758 | 103.8 | 328 | 35483 | 108.2 | 1.3 | 1.2 | 1.0 |
| Cff062; BRIP 70610 | 2018 | NSW | Mullaley | *V. radiata* | 2.0 | 0.315 | 13853 | 7826 | 97.6 | 137.3 | 0.7 | 105325 | 18906234 | 179.5 | 861 | 118118 | 137.2 | 328 | 46760 | 142.6 | 1.3 | 1.3 | 1.0 |
| Cff109 | 2018 | Qld | Brookstead | *V. radiata* | - | - | 14064 | 4202 | 99.0 | 73.7 | 1.3 | 113434 | 13041035 | 115.0 | 865 | 71383 | 82.5 | 328 | 26142 | 79.7 | 1.4 | 1.4 | 1.0 |
| Cff014; BRIP 70601 | 2018 | NSW | Mullaley | *V. radiata* | - | - | 24613 | 7077 | 173.3 | 124.2 | 1.4 | 105360 | 17880733 | 169.7 | 865 | 105344 | 121.8 | 328 | 38666 | 117.9 | 1.4 | 1.4 | 1.0 |
| Cff003; BRIP 70600 | 2018 | NSW | Mullaley | *V. radiata* | 1.9 | 0.051 | 24478 | 5917 | 172.4 | 103.8 | 1.7 | 105357 | 15494150 | 147.1 | 865 | 89969 | 104.0 | 328 | 40846 | 124.5 | 1.4 | 1.2 | 0.8 |
| Cff110; BRIP 70613 | 2018 | Qld | Brookstead | *V. radiata* | - | - | 21629 | 5521 | 152.3 | 96.9 | 1.6 | 113434 | 16653269 | 146.8 | 865 | 87297 | 100.9 | 328 | 33912 | 103.4 | 1.5 | 1.4 | 1.0 |
| Cff020 | 2018 | NSW | Mullaley | *V. radiata* | - | - | 15162 | 4526 | 106.8 | 79.4 | 1.3 | 105087 | 11864749 | 112.9 | 865 | 66484 | 76.9 | 328 | 22202 | 67.7 | 1.5 | 1.7 | 1.1 |
| CffT13694; BRIP 70617 | 2014 | Qld | Tuckerang | *V. radiata* | 1.7 | 0.061 | 25404 | 7048 | 178.9 | 123.6 | 1.4 | 113440 | 17635571 | 155.5 | 865 | 90106 | 104.2 | 328 | 42049 | 128.2 | 1.5 | 1.2 | 0.8 |
| Cff010 | 2018 | NSW | Mullaley | *V. radiata* | - | - | 32552 | 7730 | 229.2 | 135.6 | 1.7 | 113434 | 22541736 | 198.7 | 865 | 114767 | 132.7 | 328 | 40470 | 123.4 | 1.5 | 1.6 | 1.1 |
| CffT14103 | 2015 | Qld | Home Hill | *V. radiata* | - | - | 27927 | 6210 | 196.7 | 108.9 | 1.8 | 113437 | 29055435 | 256.1 | 865 | 147260 | 170.2 | 328 | 61151 | 186.4 | 1.5 | 1.4 | 0.9 |
| Cff102 | 2018 | Qld | Brookstead | *V. radiata* | - | - | 7969 | 5494 | 56.1 | 96.4 | 0.6 | 113436 | 14520611 | 128.0 | 865 | 73098 | 84.5 | 328 | 32110 | 97.9 | 1.5 | 1.3 | 0.9 |
| CffT13799A | 2014 | Qld | Warwick | *V. radiata* | 1.6 | 0.036 | 35612 | 10816 | 250.8 | 189.8 | 1.3 | 113440 | 30328117 | 267.3 | 865 | 150070 | 173.5 | 328 | 60891 | 185.6 | 1.5 | 1.4 | 0.9 |

| Isolate | Year | State | Location | Host | Estimated plasmid copies (ddPCR) | SE (ddPCR) | Total read depth across plasmid *Cff*_141 PCR amplicon (142bp) | Total read depth across *Curto*_57_*gyrB* PCR amplicon (57bp) | Mean read depth across plasmid *Cff*_141 PCR amplicon | Mean read depth across *Curto*_57_*gyrB* PCR amplicon | Relative read depth of plasmid to *gyrB* amplicon | Length of *Cff* plasmid (bp) | Total mapped read depth across *Cff* plasmid | Mean mapped read depth across *Cff* plasmid | Length of partial *gyrB* (bp) | Total mapped read depth across *gyrB* gene | Mean read depth across *gyrB* gene | Length of *rpoB* (bp) | Total mapped read depth across *rpoB* gene | Mean read depth across *rpoB* gene | Relative read depth of *Cff* plasmid to *gyrB* | Relative read depth of *Cff* plasmid to *rpoB* | Relative read depth of *gyrB* to *rpoB* |
|---|---|---|---|---|---|---|---|---|---|---|---|---|---|---|---|---|---|---|---|---|---|---|---|
| CffDA1121; BRIP 70625 | 2019 | Qld | Formartin | *V. radiata* | - | - | 25160 | 5147 | 177.2 | 90.3 | 2.0 | 113440 | 16854715 | 148.6 | 865 | 81605 | 94.3 | 328 | 36782 | 112.1 | 1.6 | 1.3 | 0.8 |
| Cff107; BRIP 70612 | 2018 | Qld | Brookstead | *V. radiata* | - | - | 25981 | 6333 | 183.0 | 111.1 | 1.6 | 113434 | 21266483 | 187.5 | 865 | 101359 | 117.2 | 328 | 37820 | 115.3 | 1.6 | 1.6 | 1.0 |
| Cff106 | 2018 | Qld | Brookstead | *V. radiata* | - | - | 43677 | 10740 | 307.6 | 188.4 | 1.6 | 113437 | 35618255 | 314.0 | 865 | 167369 | 193.5 | 328 | 61738 | 188.2 | 1.6 | 1.7 | 1.0 |
| Cff063 | 2018 | NSW | Mullaley | *V. radiata* | - | - | 29239 | 6685 | 205.9 | 117.3 | 1.8 | 105325 | 21132300 | 200.6 | 861 | 98128 | 114.0 | 328 | 41363 | 126.1 | 1.8 | 1.6 | 0.9 |
| CffDA493 | 2019 | NSW | Mullaley | *V. radiata* | - | - | 38065 | 8566 | 268.1 | 150.3 | 1.8 | 113436 | 29825111 | 262.9 | 865 | 129004 | 149.1 | 328 | 52738 | 160.8 | 1.8 | 1.6 | 0.9 |
| CffT14074 | 2015 | Qld | Emerald | *V. radiata* | - | - | 50783 | 10750 | 357.6 | 188.6 | 1.9 | 113436 | 38926823 | 343.2 | 865 | 167184 | 193.3 | 328 | 58299 | 177.7 | 1.8 | 1.9 | 1.1 |
| CffT14089; BRIP 70621 | 2015 | Qld | Biloela | *V. radiata* | 2.5 | 0.071 | 60532 | 13627 | 426.3 | 239.1 | 1.8 | 113437 | 43722022 | 385.4 | 865 | 186813 | 216.0 | 328 | 72293 | 220.4 | 1.8 | 1.7 | 1.0 |
| CffDA1155 | 2019 | Qld | Bongeen | *V. radiata* | - | - | 22636 | 4824 | 159.4 | 84.6 | 1.9 | 113435 | 17372042 | 153.1 | 865 | 73577 | 85.1 | 328 | 23491 | 71.6 | 1.8 | 2.1 | 1.2 |
| Cff033 | 2018 | NSW | Mullaley | *V. radiata* | - | - | 19311 | 4010 | 136.0 | 70.4 | 1.9 | 113436 | 14803287 | 130.5 | 865 | 62629 | 72.4 | 328 | 25524 | 77.8 | 1.8 | 1.7 | 0.9 |
| CffDA506 | 2019 | NSW | Nombi | *V. radiata* | - | - | 31430 | 7528 | 221.3 | 132.1 | 1.7 | 113436 | 24416769 | 215.2 | 865 | 103136 | 119.2 | 328 | 34302 | 104.6 | 1.8 | 2.1 | 1.1 |
| Cff012 | 2018 | NSW | Mullaley | *V. radiata* | - | - | 24697 | 4981 | 173.9 | 87.4 | 2.0 | 113436 | 18224592 | 160.7 | 865 | 76732 | 88.7 | 328 | 32980 | 100.5 | 1.8 | 1.6 | 0.9 |
| Cff002 | 2018 | NSW | Mullaley | *V. radiata* | - | - | 26846 | 5651 | 189.1 | 99.1 | 1.9 | 105357 | 19447409 | 184.6 | 865 | 87205 | 100.8 | 328 | 40656 | 124.0 | 1.8 | 1.5 | 0.8 |
| Cff064 | 2018 | NSW | Mullaley | *V. radiata* | 1.7 | 0.084 | 32912 | 7005 | 231.8 | 122.9 | 1.9 | 105325 | 21798481 | 207.0 | 861 | 97262 | 113.0 | 328 | 42491 | 129.5 | 1.8 | 1.6 | 0.9 |
| CffT13799B; BRIP 70618 | 2014 | Qld | Warwick | *V. radiata* | - | - | 22812 | 6176 | 160.6 | 108.4 | 1.5 | 113440 | 18712617 | 165.0 | 865 | 77133 | 89.2 | 328 | 28140 | 85.8 | 1.8 | 1.9 | 1.0 |
| Cff016; BRIP 70602 | 2018 | NSW | Mullaley | *V. radiata* | 1.8 | 0.107 | 22782 | 4386 | 160.4 | 76.9 | 2.1 | 113439 | 15732835 | 138.7 | 865 | 64306 | 74.3 | 328 | 22271 | 67.9 | 1.9 | 2.0 | 1.1 |
| CffUQ35 | 2018 | NSW | Lismore | *V. radiata* | - | - | 20080 | 3779 | 141.4 | 66.3 | 2.1 | 113440 | 13110071 | 115.6 | 865 | 53150 | 61.4 | 328 | 20229 | 61.7 | 1.9 | 1.9 | 1.0 |
| CffDA1149 | 2019 | Qld | Bongeen | *V. radiata* | - | - | 24903 | 5387 | 175.4 | 94.5 | 1.9 | 113436 | 19890372 | 175.3 | 865 | 80576 | 93.2 | 328 | 22449 | 68.4 | 1.9 | 2.6 | 1.4 |
| CffDA505 | 2019 | NSW | Nombi | *V. radiata* | - | - | 27938 | 5897 | 196.7 | 103.5 | 1.9 | 113437 | 23278879 | 205.2 | 865 | 93479 | 108.1 | 328 | 37576 | 114.6 | 1.9 | 1.8 | 0.9 |
| CffDA1196 | 2019 | Qld | Jondaryan | *V. radiata* | - | - | 20927 | 3049 | 147.4 | 53.5 | 2.8 | 113436 | 11815952 | 104.2 | 865 | 47190 | 54.6 | 328 | 18766 | 57.2 | 1.9 | 1.8 | 1.0 |
| Cff043; BRIP 70607 | 2018 | NSW | Mullaley | *V. radiata* | 1.4 | 0.041 | 28921 | 6771 | 203.7 | 118.8 | 1.7 | 113440 | 23027853 | 203.0 | 865 | 91583 | 105.9 | 328 | 33845 | 103.2 | 1.9 | 2.0 | 1.0 |
| CffUQ36 | 2018 | NSW | Lismore | *V. radiata* | - | - | 15366 | 3334 | 108.2 | 58.5 | 1.9 | 113440 | 11895157 | 104.9 | 865 | 47208 | 54.6 | 328 | 23379 | 71.3 | 1.9 | 1.5 | 0.8 |
| Cff118 | 2018 | Qld | Wooroolin | *V. radiata* | - | - | 21944 | 3345 | 154.5 | 58.7 | 2.6 | 113436 | 15463854 | 136.3 | 865 | 61351 | 70.9 | 328 | 22107 | 67.4 | 1.9 | 2.0 | 1.1 |
| Cff129 | 2018 | Qld | Wooroolin | *V. radiata* | - | - | 21861 | 4310 | 154.0 | 75.6 | 2.0 | 113434 | 14095801 | 124.3 | 865 | 55508 | 64.2 | 328 | 22945 | 70.0 | 1.9 | 1.8 | 0.9 |
| CffDA1153 | 2019 | Qld | Bongeen | *V. radiata* | - | - | 18145 | 4225 | 127.8 | 74.1 | 1.7 | 113436 | 16041984 | 141.4 | 865 | 63079 | 72.9 | 328 | 26993 | 82.3 | 1.9 | 1.7 | 0.9 |
| Cff192 | 2018 | Qld | Wooroolin | *V. radiata* | - | - | 70858 | 12465 | 499.0 | 218.7 | 2.3 | 113437 | 51204866 | 451.4 | 865 | 197937 | 228.8 | 328 | 65250 | 198.9 | 2.0 | 2.3 | 1.2 |
| Cff070 | 2018 | NSW | Mullaley | *V. radiata* | - | - | 34765 | 7075 | 244.8 | 124.1 | 2.0 | 113435 | 26029989 | 229.5 | 865 | 99658 | 115.2 | 328 | 29924 | 91.2 | 2.0 | 2.5 | 1.3 |
| Cff007 | 2018 | NSW | Mullaley | *V. radiata* | - | - | 38149 | 7147 | 268.7 | 125.4 | 2.1 | 113436 | 30032516 | 264.8 | 865 | 114910 | 132.8 | 328 | 40304 | 122.9 | 2.0 | 2.2 | 1.1 |
| Cff066 | 2018 | NSW | Mullaley | *V. radiata* | - | - | 24286 | 4342 | 171.0 | 76.2 | 2.2 | 113436 | 18926525 | 166.8 | 865 | 72261 | 83.5 | 328 | 32331 | 98.6 | 2.0 | 1.7 | 0.8 |

| Isolate | Year | State | Location | Host | Estimated plasmid copies (ddPCR) | SE (ddPCR) | Total read depth across plasmid *Cff*_141 PCR amplicon (142bp) | Total read depth across *Curto*_57_*gyrB* PCR amplicon (57bp) | Mean read depth across plasmid *Cff*_141 PCR amplicon | Mean read depth across *Curto*_57_*gyrB* PCR amplicon | Relative read depth of plasmid to *gyrB* amplicon | Length of *Cff* plasmid (bp) | Total mapped read depth across *Cff* plasmid | Mean mapped read depth across *Cff* plasmid | Length of partial *gyrB* (bp) | Total mapped read depth across *gyrB* gene | Mean read depth across *gyrB* gene | Length of *rpoB* (bp) | Total mapped read depth across *rpoB* gene | Mean read depth across *rpoB* gene | Relative read depth of *Cff* plasmid to *gyrB* | Relative read depth of *Cff* plasmid to *rpoB* | Relative read depth of *gyrB* to *rpoB* |
|---|---|---|---|---|---|---|---|---|---|---|---|---|---|---|---|---|---|---|---|---|---|---|---|
| Cff029; BRIP 70605 | 2018 | NSW | Mullaley | *V. radiata* | - | - | 20492 | 5951 | 144.3 | 104.4 | 1.4 | 113434 | 20554130 | 181.2 | 865 | 78201 | 90.4 | 328 | 31346 | 95.6 | 2.0 | 1.9 | 0.9 |
| CffDA270 | 2019 | Qld | Brookstead | *V. radiata* | 2.1 | 0.151 | 29968 | 4940 | 211.0 | 86.7 | 2.4 | 113436 | 20489738 | 180.6 | 865 | 77558 | 89.7 | 328 | 29708 | 90.6 | 2.0 | 2.0 | 1.0 |
| Cff009 | 2018 | NSW | Mullaley | *V. radiata* | - | - | 25470 | 6064 | 179.4 | 106.4 | 1.7 | 113436 | 23710023 | 209.0 | 865 | 89289 | 103.2 | 328 | 31889 | 97.2 | 2.0 | 2.1 | 1.1 |
| Cff200; BRIP 70615 | 2018 | Qld | Wooroolin | *V. radiata* | - | - | 27845 | 4466 | 196.1 | 78.4 | 2.5 | 113436 | 30783305 | 271.4 | 865 | 115023 | 133.0 | 328 | 34580 | 105.4 | 2.0 | 2.6 | 1.3 |
| CffDA1145 | 2019 | Qld | Bongeen | *V. radiata* | - | - | 22406 | 5710 | 157.8 | 100.2 | 1.6 | 113434 | 19984275 | 176.2 | 865 | 74624 | 86.3 | 328 | 29223 | 89.1 | 2.0 | 2.0 | 1.0 |
| Cff034 | 2018 | NSW | Mullaley | *V. radiata* | - | - | 30087 | 6050 | 211.9 | 106.1 | 2.0 | 113435 | 24777058 | 218.4 | 865 | 91552 | 105.8 | 328 | 37677 | 114.9 | 2.1 | 1.9 | 0.9 |
| Cff054 | 2018 | NSW | Mullaley | *V. radiata* | - | - | 43471 | 7751 | 306.1 | 136.0 | 2.3 | 113437 | 32505250 | 286.5 | 865 | 119482 | 138.1 | 328 | 45153 | 137.7 | 2.1 | 2.1 | 1.0 |
| CffDA494 | 2019 | NSW | Mullaley | *V. radiata* | - | - | 16391 | 3225 | 115.4 | 56.6 | 2.0 | 113434 | 13634688 | 120.2 | 865 | 50042 | 57.9 | 328 | 21869 | 66.7 | 2.1 | 1.8 | 0.9 |
| Cff056 | 2018 | NSW | Mullaley | *V. radiata* | - | - | 23300 | 3751 | 164.1 | 65.8 | 2.5 | 113434 | 16403398 | 144.6 | 865 | 58820 | 68.0 | 328 | 27545 | 84.0 | 2.1 | 1.7 | 0.8 |
| Cff052; BRIP 70608 | 2018 | NSW | Mullaley | *V. radiata* | 2.6 | 0.032 | 31564 | 4761 | 222.3 | 83.5 | 2.7 | 113436 | 26005464 | 229.3 | 865 | 92056 | 106.4 | 328 | 38635 | 117.8 | 2.2 | 1.9 | 0.9 |
| CffDA1156 | 2019 | Qld | Bongeen | *V. radiata* | - | - | 25313 | 3756 | 178.3 | 65.9 | 2.7 | 113436 | 16721549 | 147.4 | 865 | 58451 | 67.6 | 328 | 24698 | 75.3 | 2.2 | 2.0 | 0.9 |
| Cff058 | 2018 | NSW | Mullaley | *V. radiata* | - | - | 36159 | 7219 | 254.6 | 126.6 | 2.0 | 113436 | 27843465 | 245.5 | 865 | 97242 | 112.4 | 328 | 40369 | 123.1 | 2.2 | 2.0 | 0.9 |
| Cff037; BRIP 70606[a] | 2018 | NSW | Mullaley | *V. radiata* | 2.5 | 0.126 | 16637 | 3448 | 117.2 | 60.5 | 1.9 | 113436 | 13625974 | 120.1 | 865 | 47567 | 55.0 | 321 | 19120 | 59.6 | 2.2 | 2.0 | 0.9 |
| Cff199 | 2018 | Qld | Wooroolin | *V. radiata* | - | - | 62768 | 12066 | 442.0 | 211.7 | 2.1 | 113437 | 46896830 | 413.4 | 865 | 163110 | 188.6 | 328 | 69719 | 212.6 | 2.2 | 1.9 | 0.9 |
| Cff059 | 2018 | NSW | Mullaley | *V. radiata* | - | - | 35502 | 6372 | 250.0 | 111.8 | 2.2 | 113436 | 25361528 | 223.6 | 865 | 88198 | 102.0 | 328 | 36092 | 110.0 | 2.2 | 2.0 | 0.9 |
| CffDA272; BRIP 70622 | 2019 | Qld | Brookstead | *V. radiata* | - | - | 27996 | 4166 | 197.2 | 73.1 | 2.7 | 113436 | 18476788 | 162.9 | 865 | 62953 | 72.8 | 328 | 23964 | 73.1 | 2.2 | 2.2 | 1.0 |
| CffT13692; BRIP 70616 | 2014 | Qld | Tuckerang | *V. radiata* | 1.8 | 0.016 | 33646 | 5026 | 236.9 | 88.2 | 2.7 | 113440 | 23315625 | 205.5 | 865 | 77994 | 90.2 | 328 | 26781 | 81.6 | 2.3 | 2.5 | 1.1 |
| CffDA1143 | 2019 | Qld | Bongeen | *V. radiata* | - | - | 35381 | 5101 | 249.2 | 89.5 | 2.8 | 113436 | 23103647 | 203.7 | 865 | 76762 | 88.7 | 328 | 30278 | 92.3 | 2.3 | 2.2 | 1.0 |
| Cff061; BRIP 70609 | 2018 | NSW | Mullaley | *V. radiata* | - | - | 42559 | 6509 | 299.7 | 114.2 | 2.6 | 113436 | 31013359 | 273.4 | 865 | 101473 | 117.3 | 328 | 41617 | 126.9 | 2.3 | 2.2 | 0.9 |
| Cff111 | 2018 | Qld | Brookstead | *V. radiata* | - | - | 21511 | 4155 | 151.5 | 72.9 | 2.1 | 113436 | 19890285 | 175.3 | 865 | 64702 | 74.8 | 328 | 32393 | 98.8 | 2.3 | 1.8 | 0.8 |
| DAR 56678 | 1987 | NSW | Manilla | *Vigna unguiculata* | - | - | 45237 | 5966 | 318.6 | 104.7 | 3.0 | 113431 | 29563261 | 260.6 | 865 | 96110 | 111.1 | 328 | 31409 | 95.8 | 2.3 | 2.7 | 1.2 |
| CffDA1152 | 2019 | Qld | Bongeen | *V. radiata* | - | - | 34330 | 5231 | 241.8 | 91.8 | 2.6 | 113437 | 24978046 | 220.2 | 865 | 80103 | 92.6 | 328 | 29302 | 89.3 | 2.4 | 2.5 | 1.0 |
| Cff026 | 2018 | NSW | Mullaley | *V. radiata* | - | - | 36281 | 4972 | 255.5 | 87.2 | 2.9 | 113436 | 26007574 | 229.3 | 865 | 83287 | 96.3 | 328 | 31575 | 96.3 | 2.4 | 2.4 | 1.0 |
| CffDA492 | 2019 | Qld | Bongeen | *V. radiata* | - | - | 29032 | 4581 | 204.5 | 80.4 | 2.5 | 113436 | 21898713 | 193.0 | 865 | 69941 | 80.9 | 328 | 32281 | 98.4 | 2.4 | 2.0 | 0.8 |
| DAR 58699 | 1987 | NSW | Dubbo | *V. radiata* | - | - | 45622 | 7260 | 321.3 | 127.4 | 2.5 | 113440 | 34479832 | 303.9 | 865 | 109800 | 126.9 | 328 | 45896 | 139.9 | 2.4 | 2.2 | 0.9 |
| DAR 58702 | 1987 | NSW | Trangie | *Vigna mungo* | - | - | 49988 | 8630 | 352.0 | 151.4 | 2.3 | 113440 | 36082239 | 318.1 | 865 | 114751 | 132.7 | 328 | 39950 | 121.8 | 2.4 | 2.6 | 1.1 |
| CffUQ18 | 2016 | Qld | Kingaroy | *V. radiata* | - | - | 51869 | 7153 | 365.3 | 125.5 | 2.9 | 113440 | 38481388 | 339.2 | 865 | 122235 | 141.3 | 328 | 47436 | 144.6 | 2.4 | 2.3 | 1.0 |
| CffDA1169 | 2019 | Qld | Kingsthorpe | *V. radiata* | - | - | 26358 | 4112 | 185.6 | 72.1 | 2.6 | 113436 | 20425021 | 180.1 | 865 | 64733 | 74.8 | 328 | 25208 | 76.9 | 2.4 | 2.3 | 1.0 |

| Isolate | Year | State | Location | Host | Estimated plasmid copies (ddPCR) | SE (ddPCR) | Total read depth across plasmid *Cff*_141 PCR amplicon (142bp) | Total read depth across *Curto*_57_*gyrB* PCR amplicon (57bp) | Mean read depth across plasmid *Cff*_141 PCR amplicon | Mean read depth across *Curto*_57_*gyrB* PCR amplicon | Relative read depth of plasmid to *gyrB* amplicon | Length of *Cff* plasmid (bp) | Total mapped read depth across *Cff* plasmid | Mean mapped read depth across *Cff* plasmid | Length of partial *gyrB* (bp) | Total mapped read depth across *gyrB* gene | Mean read depth across *gyrB* gene | Length of *rpoB* (bp) | Total mapped read depth across *rpoB* gene | Mean read depth across *rpoB* gene | Relative read depth of *Cff* plasmid to *gyrB* | Relative read depth of *Cff* plasmid to *rpoB* | Relative read depth of *gyrB* to *rpoB* |
|---|---|---|---|---|---|---|---|---|---|---|---|---|---|---|---|---|---|---|---|---|---|---|---|
| CffUQ07 | 2016 | Qld | Canaga | *V. radiata* | - | - | 66828 | 10351 | 470.6 | 181.6 | 2.6 | 105353 | 43680361 | 414.6 | 865 | 148487 | 171.7 | 328 | 63011 | 192.1 | 2.4 | 2.2 | 0.9 |
| CffUQ20 | 2016 | Qld | Kingaroy | *V. radiata* | - | - | 30710 | 4274 | 216.3 | 75.0 | 2.9 | 113436 | 18956108 | 167.1 | 865 | 59844 | 69.2 | 328 | 23295 | 71.0 | 2.4 | 2.4 | 1.0 |
| Cff068 | 2018 | NSW | Mullaley | *V. radiata* | - | - | 35850 | 6048 | 252.5 | 106.1 | 2.4 | 113436 | 30746188 | 271.0 | 865 | 95552 | 110.5 | 328 | 37608 | 114.7 | 2.5 | 2.4 | 1.0 |
| Cff108 | 2018 | Qld | Brookstead | *V. radiata* | - | - | 30721 | 4340 | 216.3 | 76.1 | 2.8 | 113437 | 24785293 | 218.5 | 865 | 76841 | 88.8 | 328 | 32674 | 99.6 | 2.5 | 2.2 | 0.9 |
| DAR 56683 | 1986 | NSW | Moree | *V. radiata* | - | - | 42309 | 7811 | 298.0 | 137.0 | 2.2 | 113440 | 33857543 | 298.5 | 865 | 104766 | 121.1 | 328 | 40938 | 124.8 | 2.5 | 2.4 | 1.0 |
| DAR 58696 | 1986 | NSW | Bingara | *V. radiata* | - | - | 40140 | 6709 | 282.7 | 117.7 | 2.4 | 113440 | 27503266 | 242.4 | 865 | 84085 | 97.2 | 328 | 38010 | 115.9 | 2.5 | 2.1 | 0.8 |
| DAR 58698 | 1987 | NSW | Dubbo | *V. unguiculata* | - | - | 58587 | 8522 | 412.6 | 149.5 | 2.8 | 113440 | 42847001 | 377.7 | 865 | 130057 | 150.4 | 328 | 46673 | 142.3 | 2.5 | 2.7 | 1.1 |
| Cff071; BRIP 70611 | 2018 | NSW | Mullaley | *V. radiata* | - | - | 34479 | 5102 | 242.8 | 89.5 | 2.7 | 113435 | 33100174 | 291.8 | 865 | 100439 | 116.1 | 328 | 40557 | 123.6 | 2.5 | 2.4 | 0.9 |
| DAR 58701 | 1987 | NSW | Dubbo | *V. unguiculata* | 1.8 | 0.043 | 57889 | 8874 | 407.7 | 155.7 | 2.6 | 113431 | 42088326 | 371.0 | 865 | 127696 | 147.6 | 328 | 48701 | 148.5 | 2.5 | 2.5 | 1.0 |
| DAR 56677 | 1987 | NSW | Nemingha | *V. radiata* | 1.7 | 0.013 | 40370 | 6123 | 284.3 | 107.4 | 2.6 | 113440 | 29815623 | 262.8 | 865 | 88970 | 102.9 | 328 | 39265 | 119.7 | 2.6 | 2.2 | 0.9 |
| DAR 58704 | 1987 | NSW | Tyrie | *V. mungo* | - | - | 77850 | 11318 | 548.2 | 198.6 | 2.8 | 113431 | 61958144 | 546.2 | 865 | 184310 | 213.1 | 328 | 76527 | 233.3 | 2.6 | 2.3 | 0.9 |
| DAR 58705 | 1987 | NSW | Rawsonville | *Vigna angularis* | - | - | 68711 | 8107 | 483.9 | 142.2 | 3.4 | 113440 | 45281003 | 399.2 | 865 | 134030 | 154.9 | 328 | 53545 | 163.2 | 2.6 | 2.4 | 0.9 |
| CffDA274; BRIP 70623[a] | 2019 | Qld | Brookstead | *V. radiata* | 2.4 | 0.009 | 28580 | 4104 | 201.3 | 72.0 | 2.8 | 113434 | 20268184 | 178.7 | 865 | 59776 | 69.1 | 328 | 25381 | 77.4 | 2.6 | 2.3 | 0.9 |
| DAR 58706 | 1987 | NSW | Trangie | *V. angularis* | 1.7 | 0.012 | 62744 | 9124 | 441.9 | 160.1 | 2.8 | 113440 | 42580937 | 375.4 | 865 | 123739 | 143.1 | 328 | 47355 | 144.4 | 2.6 | 2.6 | 1.0 |
| Cff055 | 2018 | NSW | Mullaley | *V. radiata* | - | - | 45580 | 6657 | 321.0 | 116.8 | 2.7 | 113436 | 31045189 | 273.7 | 865 | 89217 | 103.1 | 328 | 31894 | 97.2 | 2.7 | 2.8 | 1.1 |
| DAR 56675 | 1987 | NSW | Bingara | *V. radiata* | - | - | 54996 | 7497 | 387.3 | 131.5 | 2.9 | 113440 | 39469532 | 347.9 | 865 | 113205 | 130.9 | 328 | 40109 | 122.3 | 2.7 | 2.8 | 1.1 |
| CffT13932[a] | 2015 | NSW | Moree | *V. radiata* | 2.7 | 0.057 | 15088 | 2430 | 106.3 | 42.6 | 2.5 | 113434 | 11303291 | 99.6 | 865 | 32316 | 37.4 | 328 | 12850 | 39.2 | 2.7 | 2.5 | 1.0 |
| CffDA1166 | 2019 | Qld | Kingsthorpe | *V. radiata* | - | - | 20766 | 2920 | 146.2 | 51.2 | 2.9 | 113434 | 17401689 | 153.4 | 865 | 49413 | 57.1 | 328 | 18089 | 55.1 | 2.7 | 2.8 | 1.0 |
| Cff040 | 2018 | NSW | Mullaley | *V. radiata* | - | - | 39762 | 6473 | 280.0 | 113.6 | 2.5 | 113436 | 33302542 | 293.6 | 865 | 94434 | 109.2 | 321 | 37229 | 116.0 | 2.7 | 2.5 | 0.9 |
| CffDA1150 | 2019 | Qld | Bongeen | *V. radiata* | - | - | 51361 | 6219 | 361.7 | 109.1 | 3.3 | 113436 | 32912065 | 290.1 | 865 | 93325 | 107.9 | 328 | 35568 | 108.4 | 2.7 | 2.7 | 1.0 |
| CffUQ37 | 2018 | NSW | Lismore | *V. radiata* | - | - | 19692 | 3771 | 138.7 | 66.2 | 2.1 | 113440 | 14706250 | 129.6 | 865 | 41504 | 48.0 | 328 | 20621 | 62.9 | 2.7 | 2.1 | 0.8 |
| CffDA271 | 2019 | Qld | Brookstead | *V. radiata* | 1.8 | 0.188 | 68617 | 9444 | 483.2 | 165.7 | 2.9 | 113436 | 48388500 | 426.6 | 865 | 135598 | 156.8 | 328 | 47524 | 144.9 | 2.7 | 2.9 | 1.1 |
| DAR 58703 | 1987 | NSW | Rawsonville | *V. unguiculata* | 1.7 | 0.013 | 45870 | 6711 | 323.0 | 117.7 | 2.7 | 113430 | 35620654 | 314.0 | 865 | 99239 | 114.7 | 328 | 51604 | 157.3 | 2.7 | 2.0 | 0.7 |
| DAR 56686 | 1986 | NSW | Tamworth | *V. unguiculata* | - | - | 47932 | 6716 | 337.5 | 117.8 | 2.9 | 113440 | 37195212 | 327.9 | 865 | 101617 | 117.5 | 328 | 44679 | 136.2 | 2.8 | 2.4 | 0.9 |
| Cff072 | 2018 | NSW | Mullaley | *V. radiata* | - | - | 28293 | 3954 | 199.2 | 69.4 | 2.9 | 113436 | 31681102 | 279.3 | 865 | 85816 | 99.2 | 328 | 37027 | 112.9 | 2.8 | 2.5 | 0.9 |
| Cff051 | 2018 | NSW | Mullaley | *V. radiata* | - | - | 23667 | 3308 | 166.7 | 58.0 | 2.9 | 113433 | 19595131 | 172.7 | 865 | 52737 | 61.0 | 328 | 26503 | 80.8 | 2.8 | 2.1 | 0.8 |
| Cff101 | 2018 | Qld | Brookstead | *V. radiata* | - | - | 52203 | 8092 | 367.6 | 142.0 | 2.6 | 113436 | 44445834 | 391.8 | 865 | 119536 | 138.2 | 328 | 53184 | 162.1 | 2.8 | 2.4 | 0.9 |
| Cff035 | 2018 | NSW | Mullaley | *V. radiata* | - | - | 41307 | 5007 | 290.9 | 87.8 | 3.3 | 113436 | 27598748 | 243.3 | 865 | 73028 | 84.4 | 328 | 30691 | 93.6 | 2.9 | 2.6 | 0.9 |

| Isolate | Year | State | Location | Host | Estimated plasmid copies (ddPCR) | SE (ddPCR) | Total read depth across plasmid *Cff*_141 PCR amplicon (142bp) | Total read depth across *Curto*_57_*gyrB* PCR amplicon (57bp) | Mean read depth across plasmid *Cff*_141 PCR amplicon | Mean read depth across *Curto*_57_*gyrB* PCR amplicon | Relative read depth of plasmid to *gyrB* amplicon | Length of *Cff* plasmid (bp) | Total mapped read depth across *Cff* plasmid | Mean mapped read depth across *Cff* plasmid | Length of partial *gyrB* (bp) | Total mapped read depth across *gyrB* gene | Mean read depth across *gyrB* gene | Length of *rpoB* (bp) | Total mapped read depth across *rpoB* gene | Mean read depth across *rpoB* gene | Relative read depth of *Cff* plasmid to *gyrB* | Relative read depth of *Cff* plasmid to *rpoB* | Relative read depth of *gyrB* to *rpoB* |
|---|---|---|---|---|---|---|---|---|---|---|---|---|---|---|---|---|---|---|---|---|---|---|---|
| Cff069 | 2018 | NSW | Mullaley | *V. radiata* | - | - | 48462 | 5802 | 341.3 | 101.8 | 3.4 | 113437 | 32038134 | 282.4 | 865 | 83263 | 96.3 | 328 | 30143 | 91.9 | 2.9 | 3.1 | 1.0 |
| DAR 56684 | 1986 | NSW | Boggabri | *V. radiata* | - | - | 54299 | 7776 | 382.4 | 136.4 | 2.8 | 113440 | 44464407 | 392.0 | 865 | 115297 | 133.3 | 328 | 44609 | 136.0 | 2.9 | 2.9 | 1.0 |
| Cff018 | 2018 | NSW | Mullaley | *V. radiata* | - | - | 27437 | 3475 | 193.2 | 61.0 | 3.2 | 113436 | 18724764 | 165.1 | 865 | 47695 | 55.1 | 328 | 20610 | 62.8 | 3.0 | 2.6 | 0.9 |
| CffUQ26 | 2018 | Qld | Warwick | *V. radiata* | - | - | 108210 | 14822 | 762.0 | 260.0 | 2.9 | 113437 | 84573117 | 745.6 | 865 | 200462 | 231.7 | 328 | 76228 | 232.4 | 3.2 | 3.2 | 1.0 |
| CffT14011 | 2015 | Qld | Surat | *V. radiata* | - | - | 30059 | 2745 | 211.7 | 48.2 | 4.4 | 113436 | 17457669 | 153.9 | 865 | 40776 | 47.1 | 328 | 12750 | 38.9 | 3.3 | 4.0 | 1.2 |
| DAR 56685 | 1986 | NSW | Bingara | *V. radiata* | - | - | 97101 | 10464 | 683.8 | 183.6 | 3.7 | 113440 | 66389581 | 585.2 | 865 | 141762 | 163.9 | 328 | 58832 | 179.4 | 3.6 | 3.3 | 0.9 |
| DAR 56673 | 1986 | Qld | Indooroopilly | *V. radiata* | - | - | 41643 | 4638 | 293.3 | 81.4 | 3.6 | 113439 | 29797052 | 262.7 | 865 | 63301 | 73.2 | 328 | 28690 | 87.5 | 3.6 | 3.0 | 0.8 |
| DAR 58710 | 1987 | NSW | Tamworth | *Phaseolus vulgaris* | 2.0 | 0.110 | 82992 | 8304 | 584.5 | 145.7 | 4.0 | 113440 | 59038213 | 520.4 | 865 | 123632 | 142.9 | 328 | 57954 | 176.7 | 3.6 | 2.9 | 0.8 |
| DAR 56672[a] | 1986 | Qld | Biloela | *V. radiata* | 2.7 | 0.020 | 37632 | 3198 | 265.0 | 56.1 | 4.7 | 113439 | 24537757 | 216.3 | 865 | 51379 | 59.4 | 328 | 19882 | 60.6 | 3.6 | 3.6 | 1.0 |
| DAR 58697 | 1986 | NSW | Bingara | *V. radiata* | - | - | 87479 | 6928 | 616.0 | 121.5 | 5.1 | 113440 | 57891211 | 510.3 | 865 | 106578 | 123.2 | 328 | 44696 | 136.3 | 4.1 | 3.7 | 0.9 |
| CffT14141 | 2016 | Qld | Jimbour | *V. radiata* | - | - | 46864 | 2180 | 330.0 | 38.2 | 8.6 | 113436 | 28110780 | 247.8 | 865 | 36269 | 41.9 | 328 | 20725 | 63.2 | 5.9 | 3.9 | 0.7 |
| Cff089[a] | 2018 | Qld | Brookstead | *V. radiata* | 0.0 | 0 | 0 | 4473 | 0.0 | 78.5 | na | 0 | 0 | 0.0 | 865 | 72424 | 83.7 | 328 | 31696 | 96.6 | na | na | 0.9 |
| CffDA535; BRIP 70624 | 2019 | NSW | Sherwood | *Glycine max* | 0.0 | 0 | 0 | 11313 | 0.0 | 198.5 | na | 0 | 0 | 0.0 | 865 | 149460 | 172.8 | 328 | 63385 | 193.2 | na | na | 0.9 |
| Cff080 | 2018 | Qld | Brookstead | *V. radiata* | - | - | 0 | 7317 | 0.0 | 128.4 | na | 0 | 0 | 0.0 | 865 | 99025 | 114.5 | 328 | 43609 | 133.0 | na | na | 0.9 |
| CffDA027 | unknown | Qld | Missen Flat | *V. radiata* | - | - | 0 | 0 | 0.0 | 0.0 | na | 0 | 0 | 0.0 | 225 | 12542 | 92.2 | 216 | 19482 | 90.2 | na | na | 1.0 |
| CffUQ19 | 2016 | Qld | Kingaroy | *V. radiata* | - | - | 0 | 2441 | 0.0 | 42.8 | na | 0 | 0 | 0.0 | 410 | 15679 | 141.3 | 269 | 15913 | 59.2 | na | na | 2.4 |
| DAR 58700 | 1987 | NSW | Dubbo | *G. max* | - | - | 0 | 7456 | 0.0 | 130.8 | na | 0 | 0 | 0.0 | 865 | 120190 | 138.9 | 328 | 50761 | 154.8 | na | na | 0.9 |

**Table S2** Summary of statistics for the plasmid copy number of 25 *Curtobacterium flaccumfaciens pv. flaccumfaciens* isolates.

| Statistical test | Statistic | Value |
|---|---|---|
| Wilcoxon rank sum test with continuity correction[a] | W | 1574 |
| | p-value | 0.0257 |
| Shapiro-Wilk normality test[b] | W | 0.85377 |
| | p-value | 1.64E-08 |
| Levene's Test for Homogeneity of Variance[c] | df - group | 1 |
| | df - residual | 98 |
| | F value | 0.544 |
| | Pr(>F) | 0.4625 |
| Kruskal-Wallis rank sum test[d] | chi-squared | 82.622 |
| | df | 24 |
| | p-value | 2.31E-08 |

[a] Test statistic (W) for the Wilcoxon rank-sum test for replicated experiments. The p-value represents the probability of obtaining results at least as extreme as the observed results, assuming that the null hypothesis is true.

[b] Test statistic (W) measures how well the data conforms to a normal distribution. The p-value determines the significance of the test for normality.

[c] Degree of freedom (df) associated with the group variable and the residuals. F value compares the variance between groups to the variance within groups. Pr(>F) indicates the probability that the observed differences in variances occurring by chance.

[d] Chi-squared statistic measures the differences among the rank sums of the groups. Degree of freedom (df) calculated as the number of groups minus one. The p-value is the probability of observing a chi-squared statistic at least as extreme as the observed results, assuming that the null hypothesis is true.

**Table S3** Summary of statistics for visual disease symptoms at 15 days after inoculation of mung bean plants (cv. Opal-AU) with six *Curotobacterium flaccumfaciens* pv. *flaccumfaciens* isolates and a water control treatment.

| Statistical test | Statistic | Value |
|---|---|---|
| Wilcoxon rank sum test with continuity correction[a] | W | 413.5 |
| | p-value | 0.7276 |
| Shapiro-Wilk normality test[b] | W | 0.88737 |
| | p-value | 8.11E-05 |
| Levene's Test for Homogeneity of Variance[c] | df - group | 1 |
| | df - residual | 54 |
| | F value | 0.0094 |
| | Pr(>F) | 0.9233 |
| Kruskal-Wallis rank sum test[d] | chi-squared | 45.215 |
| | df | 6 |
| | p-value | 4.24E-08 |

[a] Test statistic (W) for the Wilcoxon rank-sum test for replicated experiments. The p-value represents the probability of obtaining results at least as extreme as the observed results, assuming that the null hypothesis is true.

[b] Test statistic (W) measures how well the data conforms to a normal distribution. The p-value determines the significance of the test for normality.

[c] Degree of freedom (df) associated with the group variable and the residuals. F value compares the variance between groups to the variance within groups. Pr(>F) indicates the probability that the observed differences in variances occurring by chance.

[d] Chi-squared statistic measures the differences among the rank sums of the groups. Degree of freedom (df) calculated as the number of groups minus one. The p-value is the probability of observing a chi-squared statistic at least as extreme as the observed results, assuming that the null hypothesis is true.

**Table S4** Summary of statistics for trifoliate leaf dry weights at 15 days after inoculation of mung bean plants (cv. Opal-AU) with six *Curotobacterium flaccumfaciens* pv. *flaccumfaciens* isolates and a water control treatment.

| Statistical test | Statistic | Value |
|---|---|---|
| Wilcoxon rank sum test with continuity correction[a] | W | 305.5 |
| | p-value | 0.1587 |
| Shapiro-Wilk normality test[b] | W | 0.92768 |
| | p-value | 0.002397 |
| Levene's Test for Homogeneity of Variance[c] | df - group | 1 |
| | df - residual | 54 |
| | F value | 2.0071 |
| | Pr(>F) | 0.1623 |
| Kruskal-Wallis rank sum test[d] | chi-squared | 22.333 |
| | df | 6 |
| | p-value | 0.001053 |

[a] Test statistic (W) for the Wilcoxon rank-sum test for replicated experiments. The p-value represents the probability of obtaining results at least as extreme as the observed results, assuming that the null hypothesis is true.

[b] Test statistic (W) measures how well the data conforms to a normal distribution. The p-value determines the significance of the test for normality.

[c] Degree of freedom (df) associated with the group variable and the residuals. F value compares the variance between groups to the variance within groups. Pr(>F) indicates the probability that the observed differences in variances occurring by chance.

[d] Chi-squared statistic measures the differences among the rank sums of the groups. Degree of freedom (df) calculated as the number of groups minus one. The p-value is the probability of observing a chi-squared statistic at least as extreme as the observed results, assuming that the null hypothesis is true.

**Table S5** Summary of statistics for *Curotobacterium flaccumfaciens* pv. *flaccumfaciens* (*Cff*) DNA (log[x+1]) in total leaf tissue samples at 15 days after inoculation of mung bean plants (cv. Opal-AU) with six *Cff* isolates and a water control treatment.

| Statistical test | Statistic | Value |
|---|---|---|
| Wilcoxon rank sum test with continuity correction[a] | W | 193.5 |
| | p-value | 0.001075 |
| Shapiro-Wilk normality test[b] | W | 0.79336 |
| | p-value | 1.94E-07 |
| Levene's Test for Homogeneity of Variance [c] | df - group | 1 |
| | df - residual | 54 |
| | F value | 0.2707 |
| | Pr(>F) | 0.605 |
| Kruskal-Wallis rank sum test[d] | chi-squared | 35.1407 |
| | df | 6 |
| | p-value | 4.05E-06 |

[a] Test statistic (W) for the Wilcoxon rank-sum test for replicated experiments. The p-value represents the probability of obtaining results at least as extreme as the observed results, assuming that the null hypothesis is true.

[b] Test statistic (W) measures how well the data conforms to a normal distribution. The p-value determines the significance of the test for normality.

[c] Degree of freedom (df) associated with the group variable and the residuals. F value compares the variance between groups to the variance within groups. Pr(>F) indicates the probability that the observed differences in variances occurring by chance.

[d] Chi-squared statistic measures the differences among the rank sums of the groups. Degree of freedom (df) calculated as the number of groups minus one. The p-value is the probability of observing a chi-squared statistic at least as extreme as the observed results, assuming that the null hypothesis is true.

**Table S6** Fungal and bacterial isolates used to provide non-target DNA for PCR assay optimisation.

| Isolate[a] | Species |
|---|---|
| BRIP 64075 i | *Alternaria* spp. |
| BRIP 65025 a | *Bipolaris sorokiniana* |
| BRIP 62449 a | *Cercospora* spp. |
| BRIP 63784 c | *Diaporthe kongii* |
| BRIP 74398 | *Pseudomonas* spp. |
| BRIP 74313 | *Pseudomonas* spp. |
| BRIP 74322 | *Pseudomonas* spp. |
| BRIP 74407 | *Pseudomonas savastanoi* pv. *phaseolicola* |
| BRIP 74320 | *Pseudomonas savastanoi* pv. *phaseolicola* |

[a] Isolates obtained from the Queensland Department of Primary Industries Plant Pathology Herbarium.

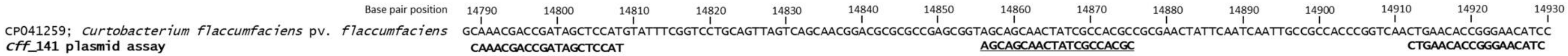


**Figure S1.** Alignment of the *Cff*_141 plasmid assay primers (bold) and probe (bold and underlined) against the plasmid sequence of *C. flaccumfaciens* pv. *flaccumfaciens* (*Cff*), partially covering a trypsin-like serine protease gene sequence. GenBank accession number CP071884 was used for base pair reference positions. This sequence was conserved amongst *C. flaccumfaciens* pv. *flaccumfaciens* isolates containing the plasmid pCff119, pCFF113 or Cff1.

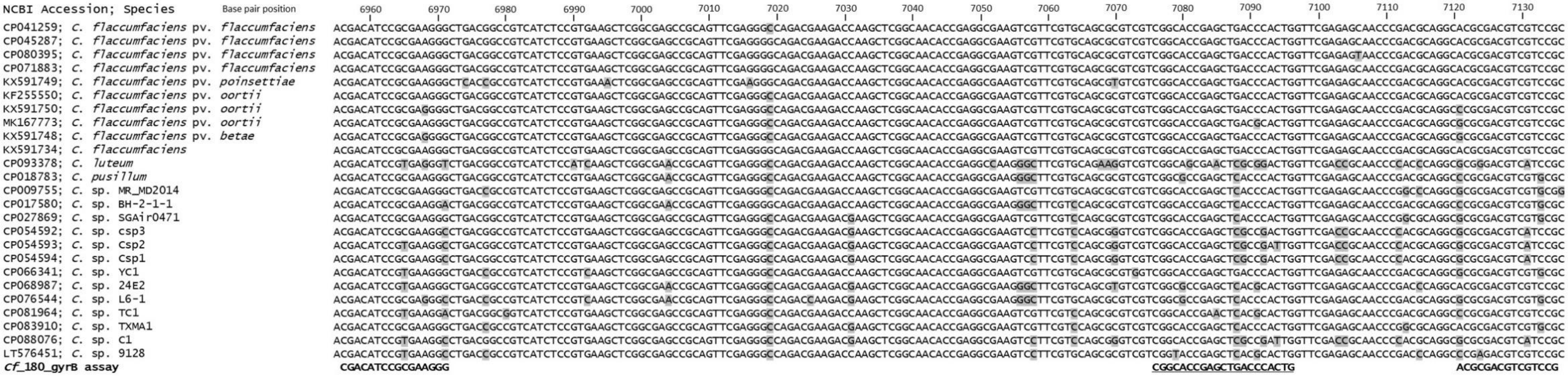


**Figure S2.** Alignment of the *Cf*_180_*gyrB* assay primers (bold) and probe (bold and underlined) against gyrase b (*gyrb*) gene sequences of a group of *Curtobacterium* species. GenBank accession number CP071883 was used for base pair reference positions. Polymorphisms among the sequences are indicated by shading (black text on grey).

| NCBI Accession; Species | Base pair position: 6780 · 6790 · 6800 · 6810 · 6820 · 6830 |
|---|---|
| CP041259; *C. flaccumfaciens* pv. *flaccumfaciens* | CGCGATGCAGTGGAACACCTCCTACCAGGAGAGCGTCCACACCTTCGCGAACACGATCA |
| CP045287; *C. flaccumfaciens* pv. *flaccumfaciens* | CGCGATGCAGTGGAACACCTCGTACCAGGAGAGCGTCCACACCTTCGCGAACACGATCA |
| CP080395; *C. flaccumfaciens* pv. *flaccumfaciens* | CGCGATGCAGTGGAACACCTCGTACCAGGAGAGCGTCCACACCTTCGCGAACACGATCA |
| KX591721; *C. flaccumfaciens* pv. *flaccumfaciens* | CGCGATGCAGTGGAACACCTCGTACCAGGAGAGCGTCCAAACCTTCGCGAACACGATCA |
| CP071883; *C. flaccumfaciens* pv. *flaccumfaciens* | CGCGATGCAGTGGAACACCTCGTACCAGGAGAGCGTCCACACGTTCGCGAACACGATCA |
| KX591749; *C. flaccumfaciens* pv. *poinsettiae* | CGCGATGCAGTGGAACACCTCGTACCAGGAGAGCGTCCACACCTTCGCGAACACGATCA |
| KX591750; *C. flaccumfaciens* pv. *oortii* | CGCGATGCAGTGGAACACCTCGTACCAGGAGAGCGTCCACACCTTCGCGAACACGATCA |
| MK167773; *C. flaccumfaciens* pv. *oortii* | CGCCATGCAGTGGAACACCTCGTACCAGGAGAGCGTCCACACCTTCGCGAACACGATCA |
| KX591748; *C. flaccumfaciens* pv. *betae* | CGCGATGCAGTGGAACACCTCGTACCAGGAGAGCGTCCACACCTTCGCGAACACGATCA |
| AM410845; *C. albidum* | CGCGATGCAGTGGAACACCTCGTACCAGGAGAGCGTCCACACCTTCGCGAACACGATCA |
| AM410844; *C. citreum* | CGCGATGCAGTGGAACACCTCGTACCAGGAGAGCGTCCACACCTTCGCGAACACGATCA |
| AM410846; *C. herbarum* | CGCGATGCAGTGGAACACCTCCTACCAGGAGAGCGTCCACACCTTCGCGAACACGATCA |
| CP093378; *C. luteum* | CGCGATGCAGTGGAACACCTCCTACCAGGAGAGCGTCCACACCTTCGCGAACACGATCA |
| CP018783; *C. pusillum* | CGCGATGCAGTGGAACACCTCCTACCAGGAGAGCGTCCACACCTTCGCGAACACGATCA |
| CP009755; *C.* sp. MR_MD2014 | CGCCATGCAGTGGAACACCTCCTACCAGGAGAGCGTCCACACCTTCGCGAACACGATCA |
| CP017580; *C.* sp. BH-2-1-1 | CGCGATGCAGTGGAACACCTCCTACCAAGAGAGCGTCCACACCTTCGCGAACACGATCA |
| CP027869; *C.* sp. SGAir0471 | CGCCATGCAGTGGAACACCTCGTACCAGGAGAGCGTCCACACCTTCGCGAACACGATCA |
| CP054592; *C.* sp. csp3 | CGCGATGCAGTGGAACACCTCGTACCAGGAGAGCGTCCACACCTTCGCGAACACGATCA |
| CP054593; *C.* sp. Csp2 | CGCGATGCAGTGGAACACCTCGTACCAGGAGAGCGTCCACACCTTCGCGAACACGATCA |
| CP054594; *C.* sp. Csp1 | CGCGATGCAGTGGAACACCTCGTACCAGGAGAGCGTCCACACCTTCGCGAACACGATCA |
| CP066341; *C.* sp. YC1 | CGCGATGCAGTGGAACACCTCGTACCAGGAGAGCGTCCACACCTTCGCGAACACGATCA |
| CP068987; *C.* sp. 24E2 | CGCGATGCAGTGGAACACCTCGTACCAGGAGAGCGTCCACACCTTCGCGAACACGATCA |
| CP076544; *C.* sp. L6-1 | CGCGATGCAGTGGAACACCTCCTACCAGGAGAGCGTCCACACCTTCGCGAACACGATCA |
| CP081964; *C.* sp. TC1 | CGCGATGCAGTGGAACACCTCGTACCAGGAGAGCGTCCACACCTTCGCGAACACGATCA |
| CP083910; *C.* sp. TXMA1 | CGCCATGCAGTGGAACACCTCGTACCAGGAGAGCGTCCACACCTTCGCGAACACGATCA |
| CP088076; *C.* sp. C1 | CGCGATGCAGTGGAACACCTCGTACCAGGAGAGCGTCCACACCTTCGCGAACACGATCA |
| LT576451; *C.* sp. 9128 | CGCGATGCAGTGGAACACCTCGTACCAGGAGAGCGTCCACACCTTCGCGAACACGATCA |
| AM410850; *Clavibacter michiganensis* subsp. *michiganensis* | CGCGATGCAGTGGACCACCTCCTACACGGAGAGCGTCCACACCTACGCGAACACCATCA |
| **Curto_57_*gyrB* assay** | **GCGATGCAGTGGAA** <u>**TACCAGGAGAGCGTCCACAC**</u> **TCGCGAACACGATC** |

**Figure S3.** Alignment of the Curto_57_*gyrB* assay primers (bold) and probe (bold and underlined) against gyrase b (*gyrb*) gene sequences of a group of *Curtobacterium* species and *Clavibacter michiganensis* subsp. *michiganensis* (related species). GenBank accession number CP071883 was used for base pair reference positions. Polymorphisms among the sequences are indicated by shading (black text on grey).

| NCBI Accession; Species | Base pair position 1370 1380 1390 1400 1410 1420 1430 1440 1450 1460 1470 1480 1490 1500 1510 1520 |
|---|---|
| XM_014656474; *V. radiata* | ACCTCTTGGACGTTTTGCTGTGAGGGACATGCGTCAAACTGTTGCTGTGGGAGTCATCAAGAACGTTGAGAAGAAGGATCCCACTGGAGCCAAGGTCACCAAGGCTGCCCAGAAGAAGAAGTGAATCGTGCTGGGTGATTCATCAGGGGATGGTTACAA |
| ***Vrr*_157_*EF1a*** **assay** | **CCTCTTGGACGTTTTGCTGT** … **<u>CAAGGTCACCAAGGCTGCCCA</u>** … **TCATCAGGGGATGGTTACA** |

**Figure S4.** Alignment of the *Vrr*_157_*EF1a* assay primers (bold) and probe (bold and underlined) against the elongation factor 1-alpha sequence of *Vigna radiata*. Uracil bases were changed to thymine for the alignment. GenBank accession number XM_014656474 was used for base pair reference positions.
